# The immunogenic potential of recurrent cancer drug resistance mutations: an *in silico* study

**DOI:** 10.1101/845784

**Authors:** Marco Punta, Victoria Jennings, Alan Melcher, Stefano Lise

**Affiliations:** Centre for Evolution and Cancer, The Institute of Cancer Research, London; Translational Immunotherapy Team, Division of Radiotherapy and Imaging, The Institute of Cancer Research, London

## Abstract

Cancer somatic mutations have been identified as a source of antigens that can be targeted by cancer immunotherapy. In this work, expanding on previous studies, we analyse the immunogenic properties of mutations that are known to drive resistance to cancer targeted-therapies. We survey a large dataset of mutations that confer resistance to different drugs and occur in numerous genes and tumour types. We show that a significant number of these mutations are predicted *in silico* to have immunogenic potential across a large proportion of the human population. Two of these mutations had previously been experimentally validated and it was confirmed that some of their associated neopeptides elicit T-cell responses *in vitro*. The identification of potent cancer-specific antigens can be instrumental for developing more effective immunotherapies. Resistance mutations, several of which are known to recur in different patients, could be of particular interest in the context of off-the-shelf precision immunotherapies such as therapeutic cancer vaccines.

## INTRODUCTION

Cancer cells express a typically aberrant protein repertoire compared to that of normal cells. These aberrations, whether functional (drivers) or non-functional (passengers), have the potential to generate peptide antigens that are not (or are only partially) subjected to central or peripheral tolerance. As such, when presented by human leukocyte antigen (HLA) complexes on the surface of cancer cells, these antigens might lead to recognition by cytotoxic T-cells and, eventually, to immuno-mediated tumor clearance^1,2^. Tumors, however, can develop sophisticated escape mechanisms^3^ and immune evasion is now recognised as one of the hallmarks of cancer^4^.

Cancer immunotherapies seek to restore the ability of the host’s immune system to recognise tumor antigens and attack the cells that express them^5^. Checkpoint blockade therapies (CBTs), in particular, act by inhibiting immune-checkpoint receptors and thus reinvigorating the cytolytic activity of the patient’s T-cell repertoire^6^. As mentioned above, cancer cells that present an aberrant peptidome are most likely to be targeted by T-cells. Despite remarkable successes, CBTs are to date approved for treatment in a limited number of solid malignancies, with only a fraction of patients responding^7^. As great efforts are being made toward improving the scope and efficacy of CBTs^8^, there is growing interest for the identification of patient’s specific, highly immunogenic antigens that could be used for more targeted treatments^9^, possibly in combination with CBTs. These include therapeutic cancer vaccines^10–12^ that can be produced *ex vivo* and delivered to the patient in the form of peptides, peptide-encoding RNA/DNA molecules or using peptide-loaded autologous dendritic cells or viruses^13^. Identification of tumor antigens that can serve for these purposes is thus a priority^14,15^.

Historically, peptides belonging to a normal cell proteome but preferentially or almost exclusively expressed in cancer cells (‘tumour-associated antigens’ or TAAs) were the first to be targeted for the clinic^11,16,17^ along with oncoviral antigens (encoded by oncogenic viruses)^18^. Although the clinical development of vaccination strategies against TAAs continues, they are now generally regarded as less-than-ideal and often weak effectors, primarily because of incomplete tumour specificity and partial central tolerance ^13,19^. Increasingly, researchers are focusing their attention on cancer-specific peptides such as those associated with passenger mutations^10,20–26^, somatic gene fusions^27^, aberrantly expressed tumor transcripts^28^ and tumor-specific alternatively spliced isoforms^29^ and post-translational modifications^30,31^.

In this study, building on previous works^32–34^, we present a comprehensive *in silico* survey of the antigenic potential of peptides associated with cancer drug resistance mutations. Resistance mutations emerge in the context of targeted therapies, which are aimed at tumors that depend for their growth on specific oncogenes^35^. This addiction makes such tumours vulnerable, at least in principle, to drugs that inhibit the relevant protein(s). Targeted therapies are available for an increasing number of haematological and solid malignancies (e.g.,^36–38^) but a significant fraction of patients either don’t respond to treatment or eventually relapse. Intrinsic (germline or somatic) and acquired (somatic) resistance is mediated by a range of different molecular mechanisms^39^. Among them is the pre-existence (possibly, if somatic, at very low allele frequencies) or the acquisition following treatment of protein-modifying mutations on the targeted oncogenes or on other genes in the same or alternative pathways^40,41^.

Resistance mutations possess a number of properties that are appealing in the context of precision immunotherapy: they are tumor-specific, thus generating neoantigens that are less likely to be subjected to central or peripheral tolerance or to elicit an autoimmune response^42^; because they drive resistance, they are expected to be expressed in therapy-resistant clones; they are usually found on oncogenes, hence making therapy-escape by the tumor through gene down-regulation harder; and, finally, several of them are known to recur in different patients (i.e., they are not patient-specific) making them potential targets for developing off-the-shelf rather than fully-personalised and potentially highly expensive precision therapies ^43^. Here, we report on 226 resistance mutations (source: COSMIC) that pertain to numerous genes, tumor types and drugs and we study their immunogenicity in relation to a set of 1,261 individuals from the 1000 Genomes project encompassing a landscape of 195 HLA-A, -B and -C class I allotypes. We show that several of these mutations generate neopeptides that are predicted *in silico* to have immunogenic potential across a significant fraction of individuals in our dataset. In the context of previous publications that showed how neopeptides from two resistance mutations (E255K in BCR-ABL1^32^ and T790M^33,34^ in EGFR) could elicit T-cell responses *in vitro*, our results support the idea that drug resistance mutations might be an important (and potentially expanding) source of tumor antigens for precision immunotherapies. In particular, this opens up the possibility of tracking the development of resistance mutations (for example, in circulating tumour DNA), whilst patients are treated on a particular drug, and using an off-the-shelf vaccine targeting the relevant neoantigen to prolong the period of clinical benefit.

## METHODS

### Mutation datasets

We download from Marty et al^44^ the following datasets: passenger mutations (1,000 in total), recurrent mutations (1,000), germline SNPs (1,000), random mutations (3,000). In Marty et al., recurrent and passenger mutations are derived from TCGA data^45^. In particular, recurrent mutations are defined as those found within a list of 200 tumour-associated genes^46^ and observed in at least 3 TCGA samples. All TCGA mutations not occurring in the list of tumour-associated genes are considered as passengers. Germline SNPs are common germline variants that are sampled from the Exome Variant Server (1,000 in total). Finally, random mutations are generated randomly in human proteins from Ensembl (release 89; GRCh37; 3,000 in total). From the initial list of 1,000 recurrent mutations, we extract those that are observed in at least 30 TCGA patients and we label them as *driver mutations* (32 in total). As explained in the next section, in the process of generating the mutation-associated neopeptides from each of these datasets, we have to discard a certain number of mutations. The final count for each set is as follows: 961 passengers, 999 recurrent (32 of which constitute our drivers list), 970 germline SNPs and 2,758 random mutations. These final lists of mutations are reported in Supplementary Table 1 together with their PMHBR score (see below).

Our resistance mutations are extracted from the CosmicResistanceMutations.tsv file that we downloaded from the COSMIC website (COSMIC version 86). From this initial list, we manually remove a few entries (COSM5855836, COSM1731743, COSM5855814, COSM3534174, COSM763) that appear to be duplicates of other entries (COSM5855837, COSM1731742, COSM5855815, COSM3534173, COSM125370, respectively), those for which information about the exact amino acid substitution is not provided in COSMIC and non-missense mutations. Overall, we obtain 226 resistance mutations (Supplementary Table 1). Note that 4 of them also appear in our list of driver mutations (NRAS Q61R and Q61K, PIK3CA E545K, BRAF V600E). Genes, tissues, tumor subtypes and drugs to which these mutations are associated are reported in Supplementary Table 2. In COSMIC, each mutation is listed as many times as the number of patients in which it has been reported in the scientific literature. Although this can provide us with valuable information on the prevalence of a mutation in patients that have been treated with a specific drug and for a specific tumor type, comparisons across different tumor types, genes and drugs are more complicated. Indeed, the number of cases reported can be influenced by several factors, including a tumor’s incidence or the time that has passed since a drug’s approval. For example the EGFR C797S mutation, a mutation of particular clinical relevance conferring resistance to the lung carcinoma third-generation EGFR inhibitor Osimertinib^47^, is currently reported in COSMIC to have occurred in 11 patients. This is much less, for example, than the 487 records for the T790M mutation. Osimertinib, however, is a relatively recent drug (FDA-approved in 2015) when compared to some of the drugs T790M confers resistance to (e.g. Gefitinib, FDA-approved in 2003). Here, notwithstanding these limitations, we make use of the number of occurrences in COSMIC to obtain at least a rough separation between rare and more frequent resistance mutations, with the latter being the ones that are more likely to be relevant in the context of off-the-shelf cancer vaccine development. In particular, we define the “resistance>1” dataset as a subset of all resistance mutations reported in COSMIC to occur in more than one patient (114 in total, Supplementary Table 1). Yet another set of resistance mutations we use is “resistance-no-BCR-ABL1”, which contains only COSMIC resistance mutations not found in BCR-ABL1 (124 in total, Supplementary Table 1). Note that although patients reported to have multiple resistance mutations might carry compound mutations that could be generating multiple-mutant neopeptides, this type of information is generally not available from COSMIC (zygosity is also for the most part unknown); as a consequence, we consider all resistance mutations as being “isolated” mutations.

### Generation of mutation-associated peptides

In order to calculate the HLA-presentation likelihood for the peptides generated by the above sets of mutations, we need to map each mutation to a protein sequence. Here, we use human protein sequences from EnsEMBL as found in the Homo_sapiens.GRCh37.pep.all.fa.gz file (downloaded from EnsEMBL and, hereafter, referred to as “EnsEMBL protein file”). For resistance mutations, we obtain from the CosmicResistanceMutations.tsv table the EnsEMBL transcript ids, all of which have a corresponding protein entry in the EnsEMBL protein file. For all other sets of mutations, we first extract the gene id from table S3 of^44^; then, we generate (Jan 2018) from the UCSC Genome Table Browser the mapping between gene ids (HGNC symbols) and canonical EnsEMBL transcripts and, additionally, from the EnsEMBL BioMart the mapping between gene ids and non-canonical EnsEMBL transcripts. Finally, given a mutation and its associated gene id, we try to map the mutation to the EnsEMBL canonical transcript sequence for that gene id. If we are not successful, we try to map the mutation to a non-canonical transcript sequence for the same gene id. If even in this second case we cannot find any appropriate mapping, we discard the mutation. For mutations that we can map to a transcript we can then find the corresponding protein sequence in the EnsEMBL protein file. According to this protocol, occasionally, two different mutations found in the same gene may end up being mapped to two different EnsEMBL transcripts and hence protein sequences. We only consider missense mutations (single amino acid substitutions); we do not consider indels. As mentioned in the previous section, the final list of mutations that we utilise from each of the above datasets following mapping to EnsEMBL transcripts is reported in Supplementary Table 1. The EnsEMBL transcripts used for genes in these datasets are shown in Supplementary Table 3.

For each mutation part of the datasets in Supplementary Table 1, we use an inhouse Python script to generate all possible peptides of length 8 to 11 that span the mutation. For mutations that don’t fall within the first 10 or last 10 positions of a transcript this means generating a total 38 peptides (or correspondingly less otherwise). A wild-type peptide associated to a specific mutant peptide is identical to the mutant peptide except for the fact that the mutated amino acid is reverted to the wild type one.

### List of individuals with known HLA allotype combinations

We obtain a list of individuals with their associated HLA class I allotypes from ftp://ftp.1000genomes.ebi.ac.uk/vol1/ftp/technical/working/20140725_hla_genotypes/20140702_hla_diversity.txt. This dataset includes 1,267 unique individuals from the 1000 Genomes Project, covering 14 populations and 4 major ancestral groups^48^. Each individual is annotated with their 6 HLA class I allotypes. However, in several cases each of the 6 allotypes is represented by multiple entries. These typing ambiguities reflect allotypes that do not differ on exons 2 and 3 of the HLA gene, that is, the exons carrying the antigen recognition sites. In these cases, we consider only the first reported entry for each allotype. We exclude individuals that have allotypes that are not well defined at the four digit level or that are not present in the NetMHCpan-4.0 library of HLA allotypes (NetMHCpan-4.0 is the method that we use for predicting HLA-presentation, see below), these are: HLA-A03:03N, HLA-B44, HLA-C15, HLA-C14XX. HLA-C0140 and those labelled “0000”. The complete list of 1,261 individuals and associated allotype combinations that we use is given in Supplementary Table 4. In the following, we refer to this as the *1000G dataset* and use it to represent the type and frequency of HLA class I combinations that we expect to find in individuals within the general population.

### HLA-presentation scores

All HLA-presentation scores that we describe in the following are defined starting from eluted ligand likelihood percentile ranks of peptides with respect to HLA allotypes; these rank scores are obtained from the NetMHCpan-4.0 prediction method^49^.

#### Best rank (BR) HLA-presentation score of a mutation

Each missense mutation is associated to a set of (maximum 38) peptides (see **Generation of mutation-associated peptides** above). For each peptide in this set, we use the program NetMHCpan-4.0^49^ to calculate the eluted ligand likelihood percentile rank and the interaction core peptide (*Icore*) with respect to all HLA allotypes observed in the 1000G dataset (see above). The elution rank takes values in the range from 0 to 100, with lower values representing higher presentation likelihoods. The *Icore* is the part of the original peptide predicted by NetMHCpan-4.0 to be located in the HLA binding site, thus the peptide most likely to interact with the T-cells. In some of the cases in which the *Icore* is shorter than the original peptide, it may not span the mutation at all and may thus be equivalent to a wild-type peptide. We define the presentation score of a mutation with respect to a specific HLA allotype as the minimum elution rank among all associated peptides (this is the same the “Best Rank” score used in^44^) excluding those with a wild-type Icore. We call this presentation score *BR*.

#### Population-wide Median Harmonic-mean Best Rank (PMHBR) score of a mutation

In Marty et al.^44^, the authors define a patient-specific presentation score for a mutation by using a harmonic mean to combine the six best rank scores of the patient’s 6 HLA allotypes (Patient Harmonic-mean Best Rank or PHBR). Unfortunately, COSMIC does not contain information about the HLA allotype combinations of the patients that develop specific resistance mutations. As a consequence, in order to provide an equal-ground comparison between all groups of mutations, we alternatively define a score that is representative of the presentation properties of a mutation across the whole population. We calculate our Population-wide Median Harmonic-mean Best Rank (PMHBR) for a mutation *m* as:

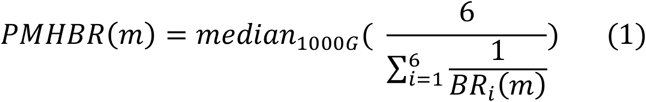

where the internal summation is taken over all 6 HLA allotypes of a given individual (2 each for HLA-A,-B and -C) and the median is taken over the 1,261 individuals from the 1000 Genomes Project for which we could obtain complete HLA class I annotation (Supplementary Table 4).

Lower PMHBR scores correspond to higher likelihoods for the mutation to be presented across the population. As remarked in Marty et al.^44^, the properties of the harmonic average imply that the lowest BRi has the biggest impact on the value of PMHBR (although all 6 terms in the summation can contribute). Because of this and the fact that HLA-C proteins are generally expressed at lower levels with respect to HLA-A and HLA-B^50^, in the supplementary materials we also report analyses in which the two HLA-C allotypes are omitted from the calculation of the harmonic average in (1).

#### Individual’s best rank

We additionally define an individual’s best rank (IBR) for a mutation *m* as the minimum BR of the mutation when considering all the HLA allotypes of the individual, that is:

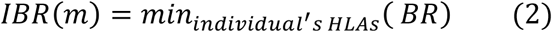

The IBR is useful for calculating the percentage of individuals in which a mutation is likely to be immunogenic according to a pre-defined threshold. For example, we can calculate the percentage of individuals for which IBR(m)<0.5 or, alternatively, <2.0 (see Results).

#### Comparison between mutant and wild-type peptide HLA-presentation scores

Given an individual with their associated HLA allotypes and a mutation, we compare the individual’s HLA-presentation scores of mutant vs wild-type peptides in the following ways. We first calculate the minimum eluted ligand likelihood percentile rank score across all of the patient’s HLA types for each pair of mutant and corresponding wild-type peptide (mutant and wild-type peptide MinRank, respectively; note that the MinRank is a property of a single peptide rather than of a mutation like the previously defined BR). We then do one of two things: (i) we ask that at least one pair exists such that the MinRank of the mutant peptide is lower than a given threshold and the MinRank of the wild-type peptide is higher than the same or different (higher) threshold or (ii) we ask that at least one pair exists such that the MinRank of the mutant peptide is lower than a given threshold and, additionally, lower than the one of the wild-type peptide. In both cases, we use thresholds of 0.5 or 2.0. Indeed, NetMHCpan-4.0 eluted ligand likelihood percentile rank score values below 0.5 are usually said to indicate high presentation likelihood, values between 0.5 and to 2.0 to indicate low presentation likelihood and values >2.0 to indicate that a peptide is not likely to be presented. We perform similar calculations for the analysis of the immunogenic potential of individual peptides in the general population.

### Statistical analysis and plots

Throughout this study, statistical analysis is performed and plots are drawn using GraphPad Prism version 8.1.1 for OS X, GraphPad Software, La Jolla California USA, www.graphpad.com. In particular, to calculate multiple comparison-adjusted p-values we perform Kruskal-Wallis tests and Dunn’s post hoc tests.

## RESULTS

In order to be immunogenic, protein peptides need to possess two fundamental properties: they have to be presentable by HLA class I complexes and they have to be able to escape central and peripheral tolerance. We start by comparing predicted HLA class I presentation scores of resistance-mutation associated neopeptides to those of neopeptides of different origin across the general population (PMHBR scores, see **Experimental Procedures**) (Figure 1 and Supplementary Figure 1 for a violin plot of the same data). We can see that passenger, germline SNP and random mutations all feature similar PMHBR score distributions. The distribution of PMHBR scores for driver mutations is instead shifted toward higher values, indicating a lower likelihood of the associated neopeptides to be HLA-presented in the general population. Although the difference between the distributions for passenger and driver mutations is not significant the trend is in line with the observation made by Marty et al. that recurrent oncogenic mutations have universally poor HLA class I presentation^44^. If we now look at resistance mutations, we see that their PMHBR scores are generally significantly lower than the ones of both passenger and driver mutations. Resistance mutation-associated neopeptides are hence predicted more likely to be HLA-presented across the general population than neopeptides generated by other types of mutations. Similar trends are observed when discarding resistance mutations reported in only one patient (the ones less relevant for off-the-shelf therapies) (Supplementary Figure 2) or when considering only HLA-A and HLA-B allotypes for computing the PMHBR score (Supplementary Figures 3 and 4). When excluding from the resistance set those mutations that occur in the BCR-ABL1 gene (constituting about half of the whole) differences appear less significant suggesting that BCR-ABL1 mutations are on average particularly immunogenic (Supplementary Figure 2 and 4). At the same time, when plotting PMHBR scores for each resistance gene separately, we see that additional genes contribute to the overall immunogenic profile of resistance mutations (e.g., ALK, MET, SMO etc.) (Supplementary Figure 5).

**Figure 1.**
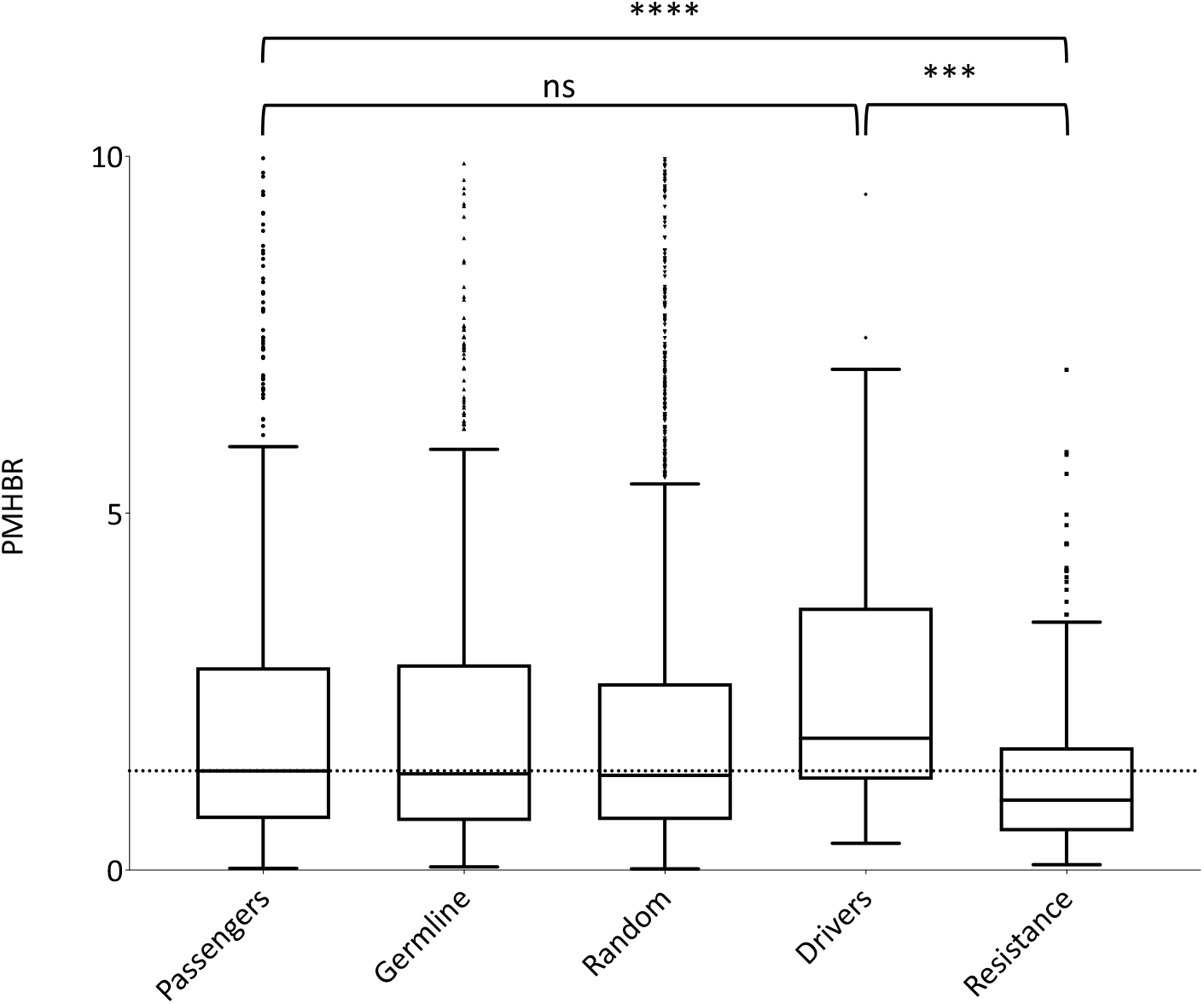
Distribution of PMHBR scores for different sets of mutations. Lower PMHBR values correspond to a higher likelihood of being presented by HLA class I complexes. The dotted horizontal line is a guide for the eye and corresponds to the value of the median of the distribution for passenger mutations. Note that for clarity the y-axis is cut at 10, thus excluding some of the distributions’ outliers. Asterisks indicate significance of pair-wise differences between PMHBR score distributions calculated using a Kruskal-Wallis test followed by Dunn’s *post hoc* test. p-values are adjusted for multiple testing (all vs all). For clarity, on the plot we report p-values for only some of the comparisons. (***) stands for p-value<0.001 and (****) stands for p-value<0.0001; (ns) stands for “not significant”. The lower and higher edges of each Tukey box represent the 25% and 75% percentile value, respectively. The horizontal line inside each box represents the median value.

In Figure 2, as examples, we show the BR score-based HLA profile (see **Experimental Procedures**) of two resistance mutations of particular clinical relevance. The EGFR C797S mutation represents a major challenge for treatment of osimertinib-resistant tumors in non-small cell lung cancer^47^. T315I, until approval of the third-generation inhibitor ponatinib^51^, was the most common mutation associated with resistance to BCR-ABL1 inhibitors^52,53^. From these profiles, we see that both mutations are predicted to generate neopeptides that produce low BR scores (that is, are predicted to have high presentation likelihood) for a wide range of common HLA allotypes. This is markedly different to what we observe for most (though not all) of the common driver mutations (see, as examples, the BR score-based HLA profiles of the two most common somatic mutations in TCGA, Supplementary Figure 6).

**Figure 2.**
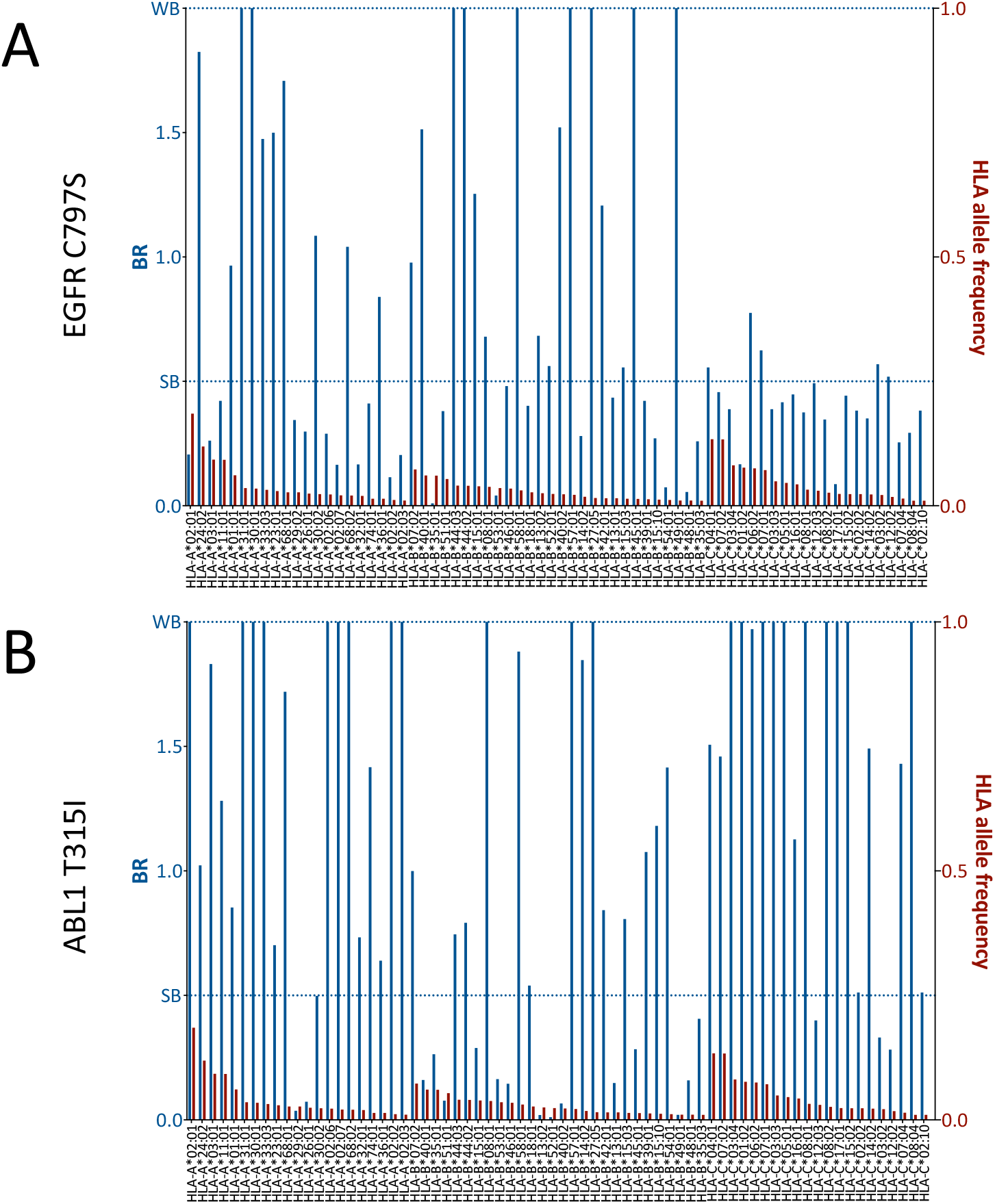
BR score-based HLA profiles of two highly clinically relevant cancer resistance mutations: **A)** C797S in EGFR and **B)** T315I in BCR-ABL1. Blue bars (primary y-axis) represent the BR scores of the mutation with respect to the HLA allotypes reported on the x-axis. Red bars (secondary y-axis) represent the frequency of each HLA allotype in the 1000G dataset. For clarity, we report BR scores for only the HLA-A, -B and -C allotypes that have frequency >1%. The two dotted lines mark elution rank value limits for strong likelihood of presentation (SB, i.e score < 0.5) and weaker likelihood of presentation (WB, i.e. 0.5<=score<2.0). For the sake of readability, we cut the primary y-axis to a value of 2.0. Note that blue bars that reach up to a value of 2.0 often correspond to BR values >2.0 and are hence cases for which no-peptide generated by the mutation is predicted likely to be presented by that specific HLA allotype.

As we mentioned in the **Introduction**, one of the most appealing characteristics of resistance mutation-associated neopeptides is that they recur in different patients. It is thus interesting to know in how many potential patients a mutation can generate neopeptides likely to be presented by HLA class I complexes. In Figure 3, for every resistance mutation recorded in at least 5 patients in COSMIC (61 mutations in total, values for all 226 mutations are in Supplementary Table 5), we calculate the percentage of individuals in our 1000G dataset that are expected to HLA-present at least one of the mutation-associated neopeptides when using an IBR score threshold of <0.5 (see also Supplementary Figure 7 for the same plot but considering only HLA-A and HLA-B allotypes for calculating the IBR score). We first notice that our results are in line with previous studies that showed that several BCR-ABL1 mutations^32^ (and T790M in EGFR^33,34^) are likely to be HLA-presented (Supplementary Figure 8), although those studies considered a much smaller set of HLA allotypes. Second and more importantly, we show that several other resistance mutations occurring in different tumour tissues (Figure 3) and in several other genes (Supplementary Figure 9) are also predicted as likely to be HLA-presented across the population. In general, we can see that 39 of 61 mutations in Figure 3 generate neopeptides that are predicted to be HLA-presented by at least 50% of individuals in our 1000G dataset. These 39 mutations occur in 6 different tumor tissues and across 12 different genes. When considering a more relaxed threshold for HLA-presentation (IBR<2.0) all but one of these mutations are predicted as “presentable” by at least half of individuals in the 1000G dataset (Supplementary Figure 10). As an example, neopeptides associated with the osimertinib resistance mutation C797S in EGFR and to T315I in BCR-ABL1 are predicted as “presentable” in 99% and 83% of individuals, respectively (for a patient-BR threshold <0.5). Interestingly, the two mutations that have been previously validated as able to elicit T-cell responses in healthy donors and patients alike (E255K in BCR-ABL1 and T790M in EGFR) are not among our top-ranked ones (21^st^ and 29^th^, respectively).

**Figure 3.**
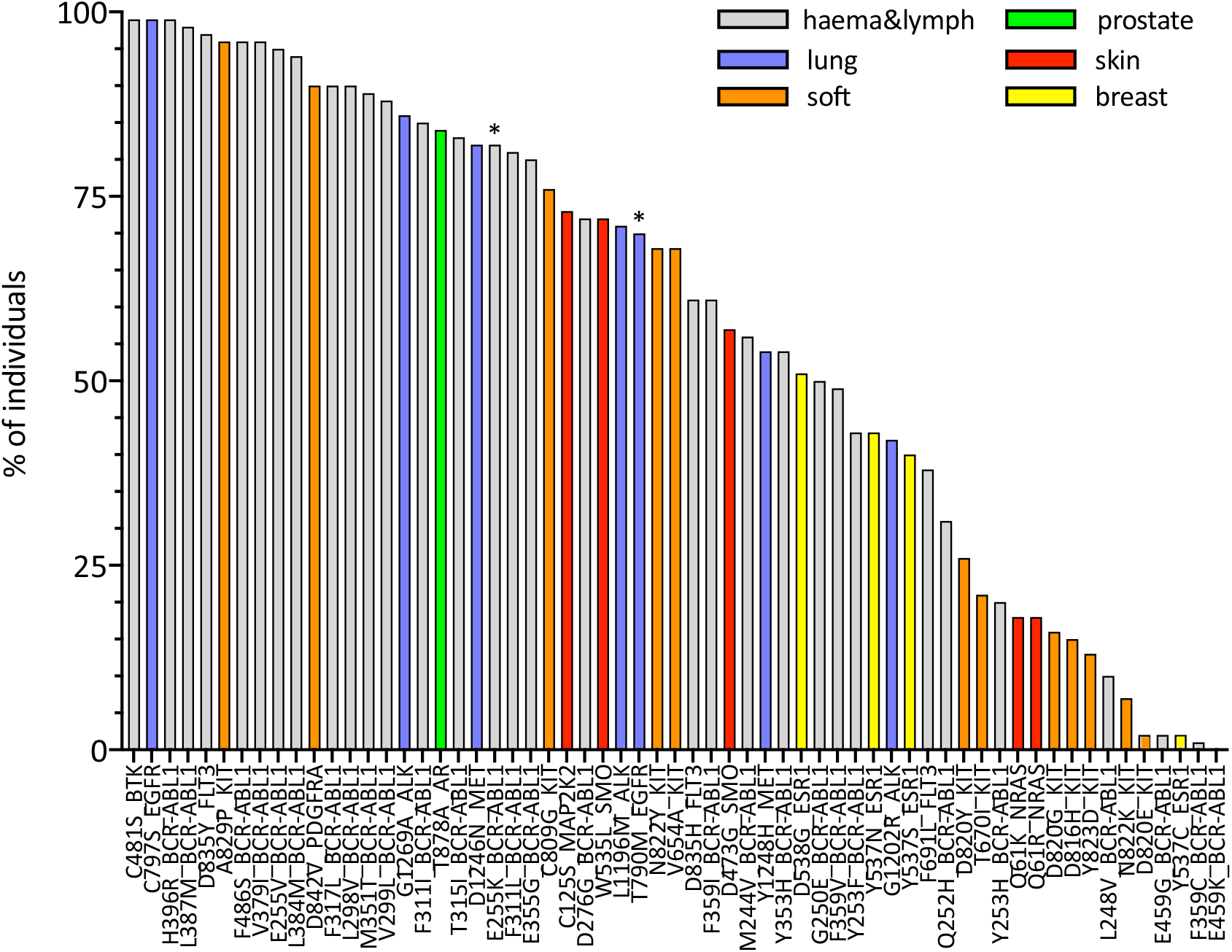
Estimates for the percentage of individuals in the general population predicted to HLA-present resistance mutation-associated neopeptides. For each mutation, the histogram illustrates the percentage of individuals in the 1000G dataset with an IBR<0.5 (the IBR score is defined in **Experimental Procedures**). Mutations on the x-axis are ordered according to decreasing percentages of individuals. For clarity, we plot only mutations that have been observed in at least 5 patients (according to COSMIC). Colours indicate the different tumour tissues in which the resistance mutations have been observed; “haema&lymph” stands for haematopoietic and lymphoid tissue. Asterisks mark mutations that have been shown to elicit T-cell responses in previous works^32–34^.

We next investigate the possibility that resistance mutation neopeptides, while likely to be HLA-presented, may be subjected to tolerance, in which case they would not be immunogenic. Under normal circumstances, tolerance ensures that there are no T-cells that can recognise germline wild type peptides, thus preventing auto-immune responses^54^. Since neopeptides are generated by somatic mutations, they are very likely to differ from any germline wild type peptide. At the same time, neopeptides originating from missense mutations, such as those that we analyse here, differ from wild type peptides only by a single amino acid substitution. Given that T-cell binding properties allow for at least some promiscuity in peptide binding affinity^55^, missense mutation-associated neopeptides might still be subjected to some degree of tolerance. A common way to identify neopeptides that are more likely to be immunogenic is to select for those that have wild type counterparts with low HLA-presentation likelihood^56^. The rationale is that if a wild type peptide is poorly presented, T-cells that bind to it and hence possibly to very similar peptides are less likely to have been negatively selected. It is important to stress, however, that even a neopeptide for which the wild type counterpart is HLA-presented may be immunogenic if the mutation it carries make it eligible to binding by a different T-cell pool with respect to the wild type. In Figure 4, we compare the presentation likelihood of resistance mutation-associated neopeptides with that of their wild type counterparts (only mutations recorded in at least 5 patients in COSMIC; values for all mutations are in Supplementary Table 5). In particular, we report the percentage of individuals in the 1000G dataset in which at least one mutant peptide is predicted highly likely to be presented (<0.5% rank) while the corresponding wild type peptide is not highly likely to be presented (>0.5% rank). We can see that for 13 mutations the percentage of individuals is at least 50% (see Supplementary Figure 11 for HLA-A and HLA-B only). Again, mutations previously shown to be immunogenic do not exhibit the highest rankings in this plot (BCR-ABL1 E255K is 11^th^ and EGFR T790M is 44^th^). In Supplementary Figures 12-15, we show the same analysis when using alternative criteria for evaluating the difference between mutant and wild type peptides. Finally, in Supplementary Figures 16 and 17, we show the resistance mutations-associated neopeptides (length 8 to 11) that we estimate to have the highest percentage of individuals more likely to present them (%rank<0.5 or %rank<2.0) than to present their wild type counterparts (%rank>0.5 or %rank>2.0 respectively). With respect to the previously validated neopeptides associated with the E255K BCR-ABL1 and T790M EGFR resistance mutations we observe the following. We predict the BCR-ABL1-associated neopeptide KVYEGVWKK to be highly likely to be HLA-presented by the HLA-A*03:01 allotype, or the allotype for which immunogenicity has been validated (%rank=0.005), and almost 100-fold more likely to be presented than its wild type counterpart EVYEGVWKK (%rank=0.33). However, since the wild type peptide is also predicted as likely to be presented (%rank<0.5), KVYEGVWKK does not fare high in our plots that use a fixed threshold for both mutant and wild type peptides while it scores definitely better when considering a <0.5% rank threshold for the mutant and simply asking that the wild type has higher ranking than the mutant peptide (Supplementary Figure 18). In contrast, we don’t predict the T790M-associated mutant peptides that have been previously validated as immunogenic (MQLMPFGCLL, LIMQLMPFGCL, IMQLMPFGC) to be likely to be presented by the experimentally validated HLA-A*02:01 allotype (%ranks=2.12, 16.0 with core peptide LIMQLMPFL and 5.74, respectively). We do however observe a separate, not previously tested T790M-associated neopeptide (LTSTVQLIM) as one that has high immunogenic potential across the population (third from top in Supplementary Figure 17).

**Figure 4.**
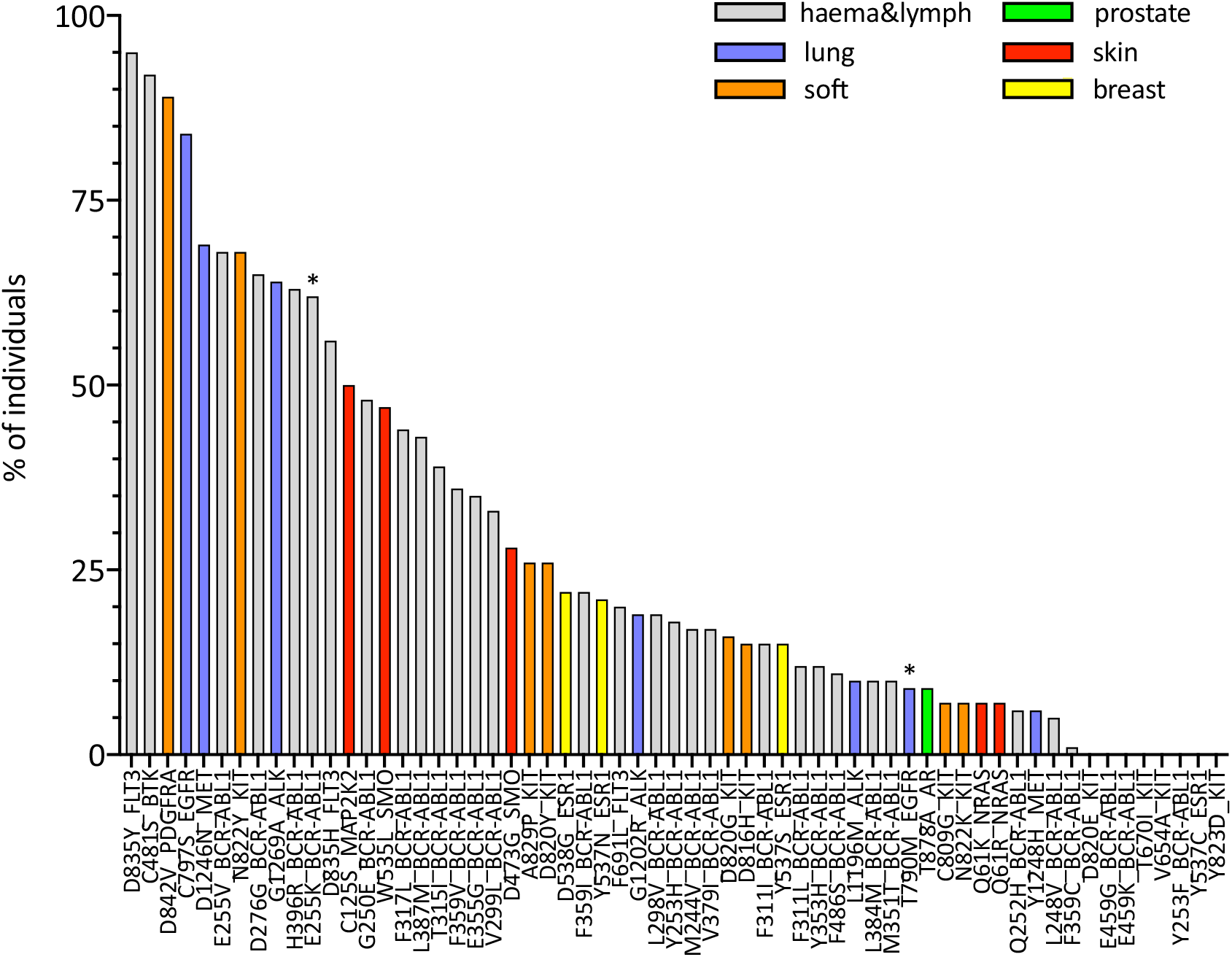
Population-wide comparison of HLA class I presentation likelihood between resistance mutation-associated mutant peptides and their corresponding wild type peptides. For each mutation, the histogram illustrates the estimated percentage of individuals for which there exists at least one mutant peptide-wild type peptide pair such that minimum eluted ligand likelihood percentile rank score across all of the individual’s HLA allotypes is <0.5 for the mutant peptide and >=0.5 for the wild type peptide. Mutations on the x-axis are ordered according to decreasing percentages of individuals. We plot only mutations that have been observed in at least 5 patients (according to COSMIC). Colours indicate the different tumour tissues in which the resistance mutations have been observed; “haema&lymph” stands for haematopoietic and lymphoid tissue. Asterisks mark mutations that have been shown to elicit T-cell responses in previous works^32–34^.

The complete list of resistance mutations that we analyse here along with estimates of the percentage of individuals in the 1000G dataset that are likely to present their associated neopeptides and, separately, that are more likely to present their associated neopeptides than their wild type counterparts can be found in Supplementary Table 5. The list of resistance mutation-associated neopeptides likely to be presented by at least 1% of individuals or, separately, more likely to presented than their wild type counterparts by at least 1% of individuals can instead be found in Supplementary Table 6 (peptides associated to resistance mutations observed in at least 5 patients, COSMIC).

## DISCUSSION AND CONCLUSIONS

Cancer immunotherapies seek to invigorate a patient’s immune response against the tumor^5^. This response is typically mediated by tumor antigens that originate from the cancer cells’ aberrant proteome. Cancer drug resistance mutations are one class of somatic aberrations that generate tumor-specific, potentially immunogenic antigens. Previous studies showed that two resistance mutations, E255K in BCR-ABL1 and T790M in EGFR, are indeed immunogenic^32–34^; additionally, Cai et al.^32^ suggested that this property may be shared by a larger number of BCR-ABL1 resistance mutations. These previous studies were based on presentability by a small number of class I HLA allotypes (5 HLA-A and 3 HLA-B allotypes in^32^ and only HLA-A*02:01 in^33,34^). We asked whether these immunogenic properties could be shared by a larger number of cancer drug resistance mutations and when considering individuals featuring a much larger number of class I HLA allotypes. Using *in silico* predictions, for the first time, we present a general survey of the immunogenicity of 226 missense resistance mutations associated with several genes (19), tissues (9) and tumor subtypes (27). We show that many of these mutations generate neopeptides that are predicted to be HLA-presented by a large proportion of the general population. Additionally, for several resistance mutations and in a significant percentage of patients, these potential neoantigens are predicted more likely to be HLA-presented than their wild type counterparts, and are therefore less likely to fall under central or peripheral tolerance. We also note that while we have considered only missense mutations, which constitute the vast majority of drug resistance mutations currently annotated in COSMIC, insertions and deletions are also known to confer resistance to some drugs. HLA-presented neopeptides generated by this type of somatic alterations would be more likely to be immunogenic as they will generally differ substantially from any wild type protein peptide^57^.

Our study comes with a number of limitations. The most obvious is that our results are based on computational predictions. Although the most recent breed of prediction methods (such as the NetMHCpan-4.0 program that we use here) integrate peptides’ HLA-elution mass spectrometry data they are still likely to over-predict the number of presented peptides^58,59^. Also, higher presentation likelihood with respect to the corresponding wild type peptide is likely to be a limited proxy for a mutant peptide’s immunogenicity (i.e., recognition by T-cells). Despite these important *caveats*, as mentioned above, computational predictions have been used previously to identify potentially immunogenic neopeptides from the BCR-ABL1 E255K and EGFR T790M resistance mutations, which were later proved effective in priming naïve T-cells^32–34^; two of these studies showed, additionally, that mature T-cells recognising these peptides could pre-exist in patients^32,33^. Further, methods that predict HLA-presentation have been widely adopted and instrumental to studies that showed neoantigen load correlation with CBT response^60^, immune-evasion by neoantigen elimination^61–63^ and investigated personalised cancer vaccines against melanoma and glioblastoma in small clinical trials^21–24^. In fact, several studies have shown that lists of predicted neoantigens are indeed enriched in neopeptides capable of stimulating T-cell responses both *in vitro* and *in vivo*^20–24^. Another potential concern is the fact that some of the proteins that develop resistance mutations to current targeted therapies are membrane-inserted. Membrane proteins are generally believed to undergo degradation in lysosomes^64^ rather than via the ubiquitin-proteasome pathway, which leads to HLA presentation of protein peptides. There is compelling evidence, however, that nevertheless membrane protein peptides are presented by HLA class I complexes^65^. Finally, although we show that many resistance mutations generate neopeptides that are predicted to be HLA-presented in most individuals, we have not ruled out the possibility that resistance mutations may be under negative selection in patients. In other words, they might occur only or primarily in those patients in which the associated neopeptides are not likely to be presented. Although negative selection has been reported for driver mutations^44^, it would seem less probable for resistance mutations which typically appear later during cancer evolution or when immune-evasion by the tumor is likely to have already occurred. One previous study, however, reported a negative correlation between response to antigens derived from the EGFR T790M mutation and the occurrence of the mutation in non-small cell lung cancer patients treated with tyrosine kinase inhibitors^33^. Unfortunately, since COSMIC does not provide us with information about the HLA allotypes of the patients in which the different resistance mutations occur, we are not able to test this hypothesis directly. If resistance mutations were under negative selection, however, it would be reasonable to expect a correlation between their observed frequency in patients and their PMHBR score (where higher PMHBR scores correspond to a lower likelihood of being presented in the general population). To reduce the impact of possible confounding factors, we consider mutations occurring on the same gene and associated with the same drug (see also **Experimental Procedures**). We take as an example the set of imatinib-associated BCR-ABL1 resistance mutations, which are the most numerous for a single drug-gene pair in COSMIC (84 total, Supplementary Table 7). In this case, we see no correlation between number of patients in which the different mutations occur and their PMHBR score (Supplementary Figure 19, differences between groups are not significant, Kruskal Wallis test). As an example, T315I or the most common imatinib-associated BCR-ABL1 mutation in COSMIC (reported in 222 patients) corresponds to a PMHBR of 0.53 compared to a median of 0.83 for all imatinib-BCR-ABL1 mutations. This appears to support the idea that at least some of these mutations might occur in patients in which they are presented and potentially immunogenic; however, a definitive answer will only come from experimental testing of the T-cell repertoire in patients carrying these mutations.

In conclusion, expanding on previous studies, we have presented data that suggests that resistance mutation-associated neoantigens could be particularly interesting targets for precision immunotherapies such as cancer vaccines^66^. Most recent work in the field has focused on tumor neoantigens associated with protein-modifying passenger mutations^21–24^. However, vaccines derived from passenger mutations, which are private, would represent fully personalised treatments with potentially high development costs and scale-up issues for translation into the clinic^43^. In contrast, recurrent neoantigens such as those potentially derived from resistance mutations could serve as a basis for developing off-the-shelf vaccines, which could be used in combination with targeted therapies, as well as with other types of immunotherapies such as CBTs. We believe that the recent advances in cancer immunotherapy and the ever-increasing number of available targeted therapies provide an unprecedented background on which to test this hypothesis.

