## Supplementary Figures for "The immunogenic potential of recurrent cancer drug resistance mutations: an *in silico* study"

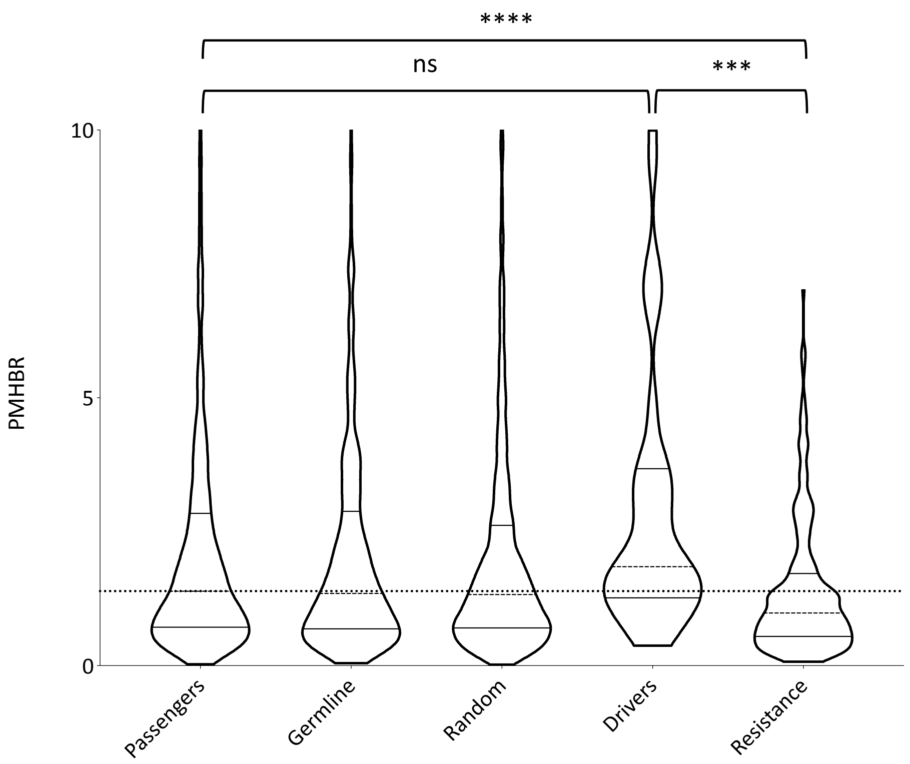


**Supplementary Figure 1.** *Same as Figure 1 but using violin plots to give a better idea of the shape of the distributions corresponding to different mutations' datasets.* Horizontal lines within each violin plot indicate 25th percentile, median (dashed) and 75th percentile (bottom, middle and top, respectively).

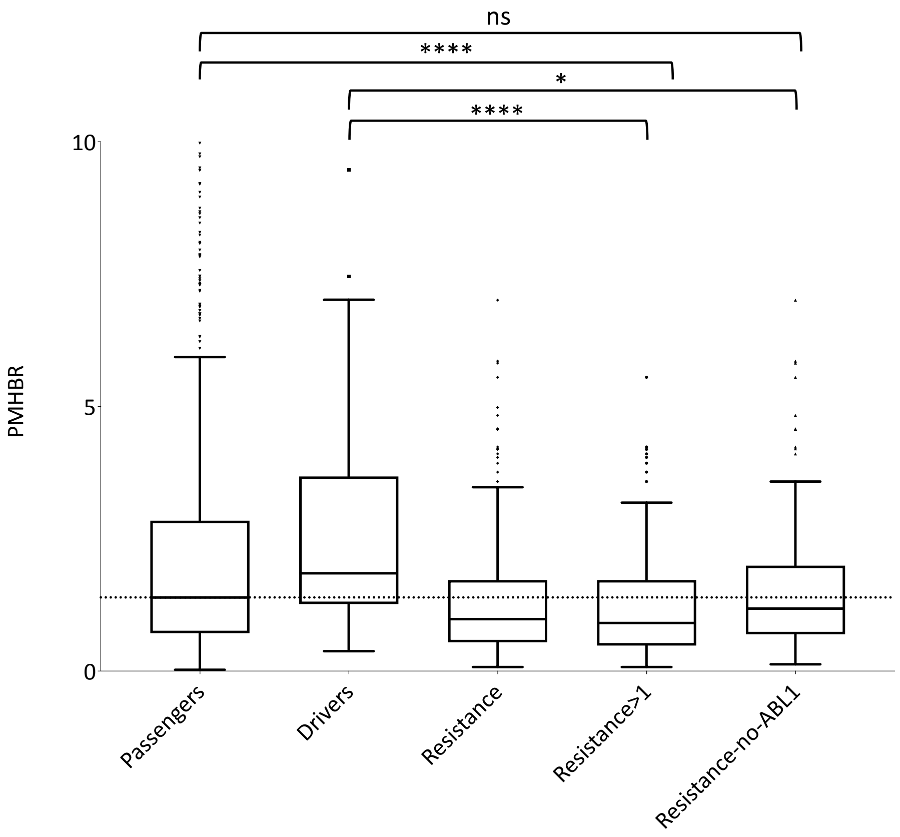


**Supplementary Figure 2.** *Similar to Figure 1 but plotting separately the PMHBR score distribution for two subsets of the resistance mutation dataset.* For a description of the different datasets see Experimental Procedures. Lower PMHBR values correspond to a higher likelihood of being presented by HLA class I complexes. The dotted horizontal line is a guide for the eye and corresponds to the value of the median of the distribution for passenger mutations. Note that for clarity the y-axis is cut at 10, thus excluding some of the distributions' outliers. Asterisks indicate significance of pair-wise differences between PMHBR score distributions calculated using a Kruskal-Wallis test followed by Dunn's *post hoc* test. p-values are adjusted for multiple testing (all vs all). For clarity, on the plot, we report p-values for only some of the comparisons. (*) stands for p-value<0.05 and (****) for p-value<0.0001; (ns) stands for "not significant". The lower and higher edges of each Tukey box represent the 25th and 75th percentile value, respectively. The horizontal line inside each box represents the median value.

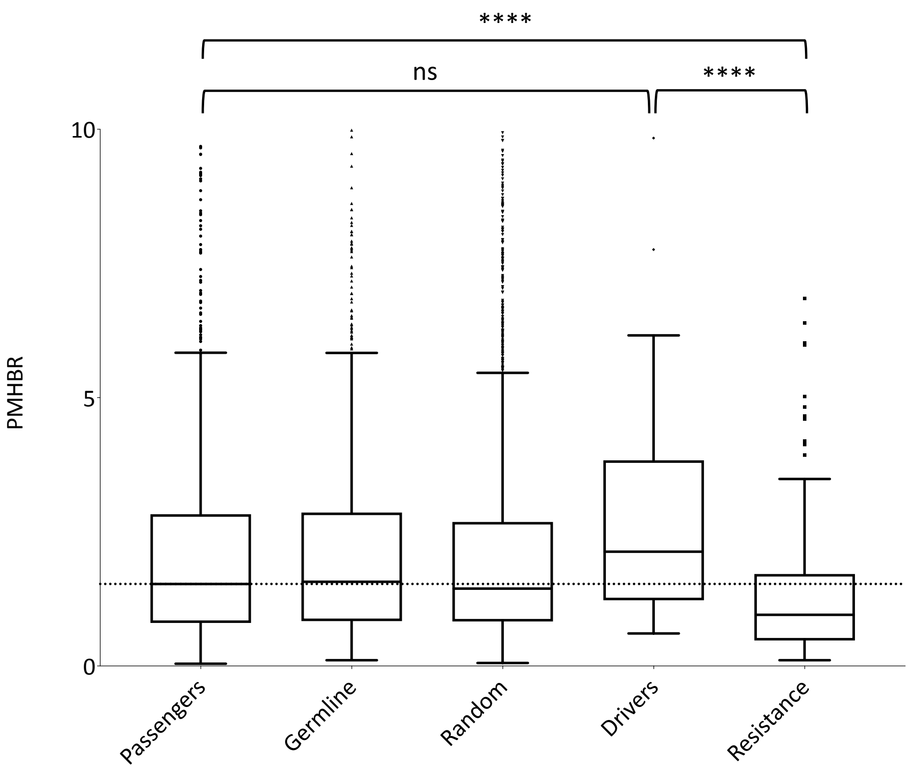


**Supplementary Figure 3.** *Equivalent to Figure 1 when using only HLA-A and HLA-B for calculating the PMHBR (Equation 1).*

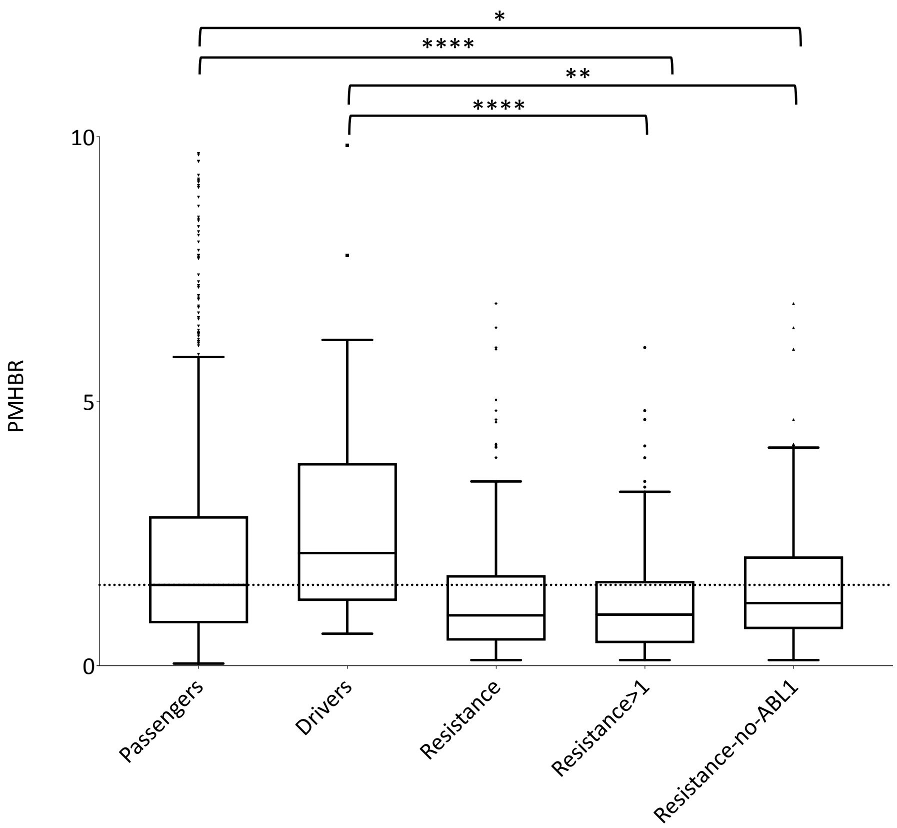


**Supplementary Figure 4.** *Equivalent to Supplementary Figure 1 when using only HLA-A and HLA-B for calculating the PMHBR (Equation 1).*

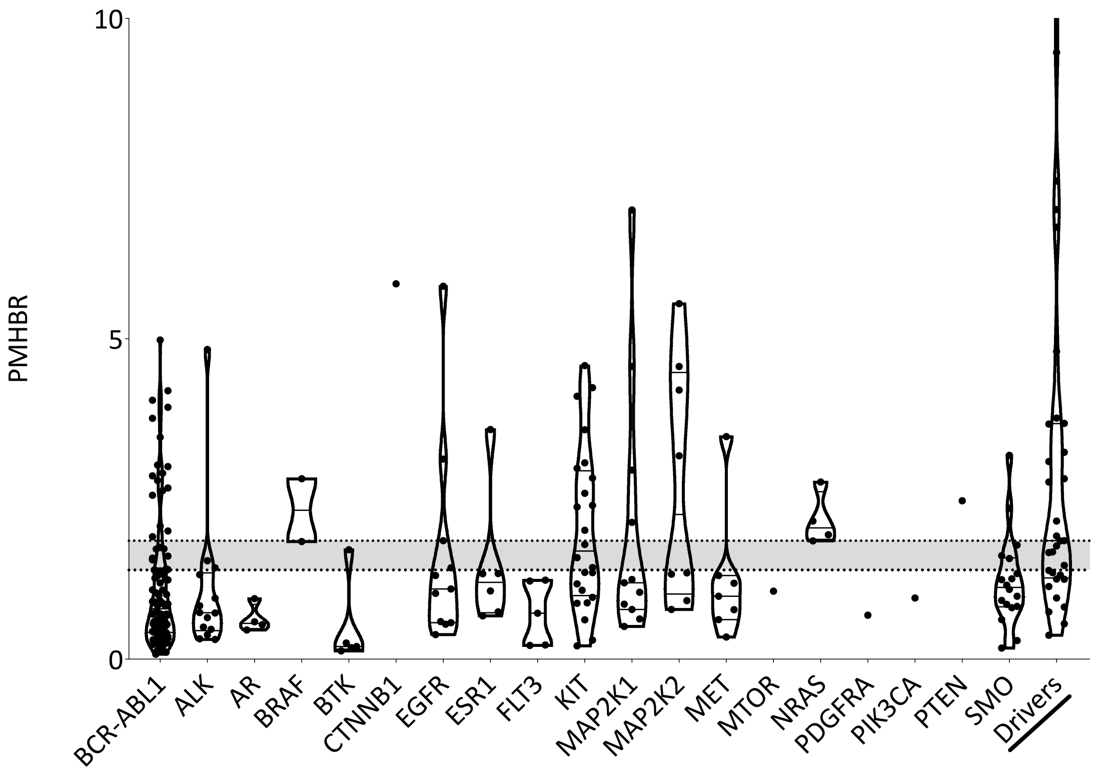


**Supplementary Figure 5.** *Violin plots of the distributions of PMHBR scores for 19 genes for which COSMIC reports resistance mutations and for driver mutations.* PMHBR scores are calculated using all 3 HLA genes (HLA-A, -B and -C). Lower PMHBR values correspond to a higher likelihood of being presented by HLA class I complexes. The dotted horizontal lines are guides for the eye and correspond to the value of the median of the distribution for passenger (bottom) and driver mutations (top), respectively (as seen for example in Figure 1). Horizontal lines within each violin plot indicate 25$^{th}$ percentile, median (dashed) and 75$^{th}$ percentile (bottom, middle and top, respectively).

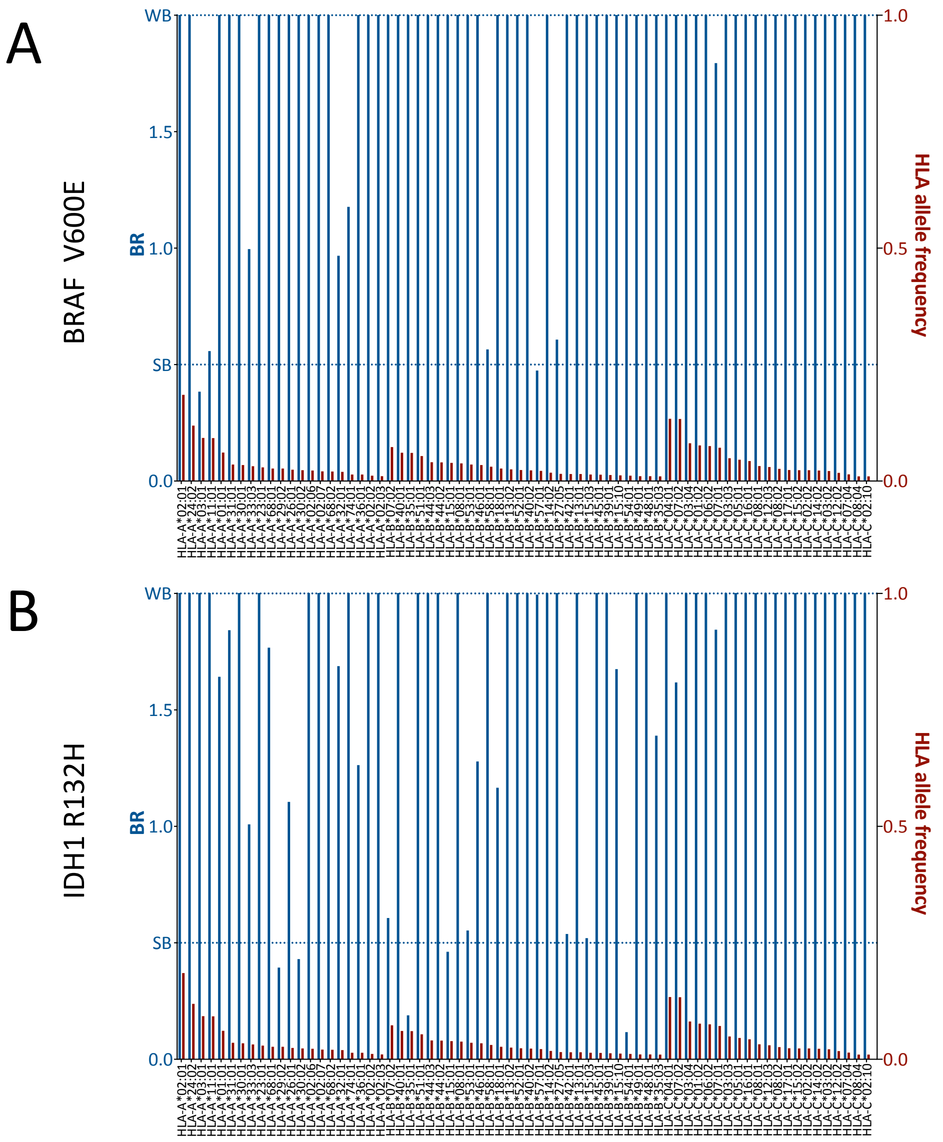


**Supplementary Figure 6.** *BR score HLA profiles of the two most common somatic mutations in TCGA patients:* ***A)*** *V600E in BRAF (present in 561 TCGA patients) and* ***B)*** *R132H in IDH1 (384 TCGA patients).* Blue bars (primary y-axis) represent the BR scores of the mutation with respect to the HLA alleles reported on the x-axis. Red bars (secondary y-axis) represent the frequency of each HLA allotype in the 1000G dataset. For clarity, we report BR scores for only the HLA-A, -B and -C allotypes that have frequency >1%. The two dotted lines mark elution rank value limits for strong likelihood of presentation (SB, i.e score < 0.5) and weaker likelihood of presentation (WB, i.e. 0.5<score<2.0). For the sake of readability, we cut the primary y-axis to a value of 2.0. Note, however, that the vast majority of BR scores (blue bars) that reach up to a value of 2.0 in the plot correspond to values >2.0 and are hence cases for which no-peptide generated by the mutation is predicted likely to be presented by that specific HLA allele.

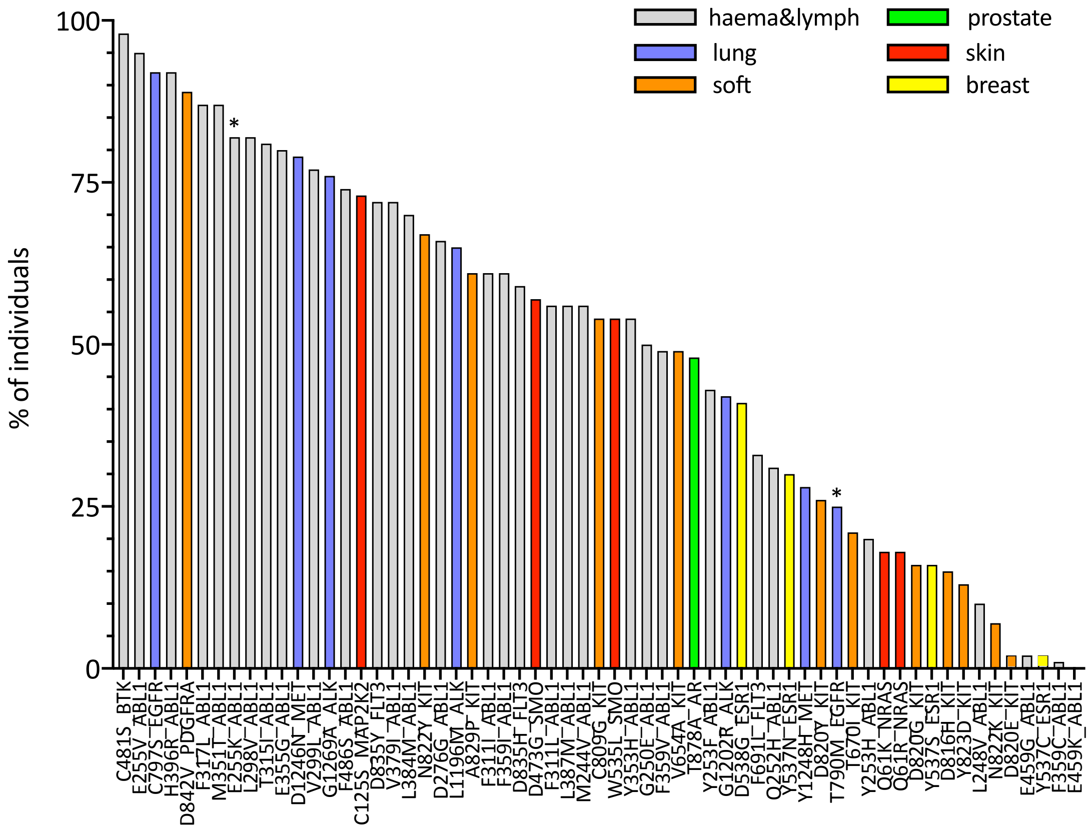


**Supplementary Figure 7.** *Equivalent to Figure 3 when using only HLA-A and HLA-B for calculating the PWHBR (Equation 1).*

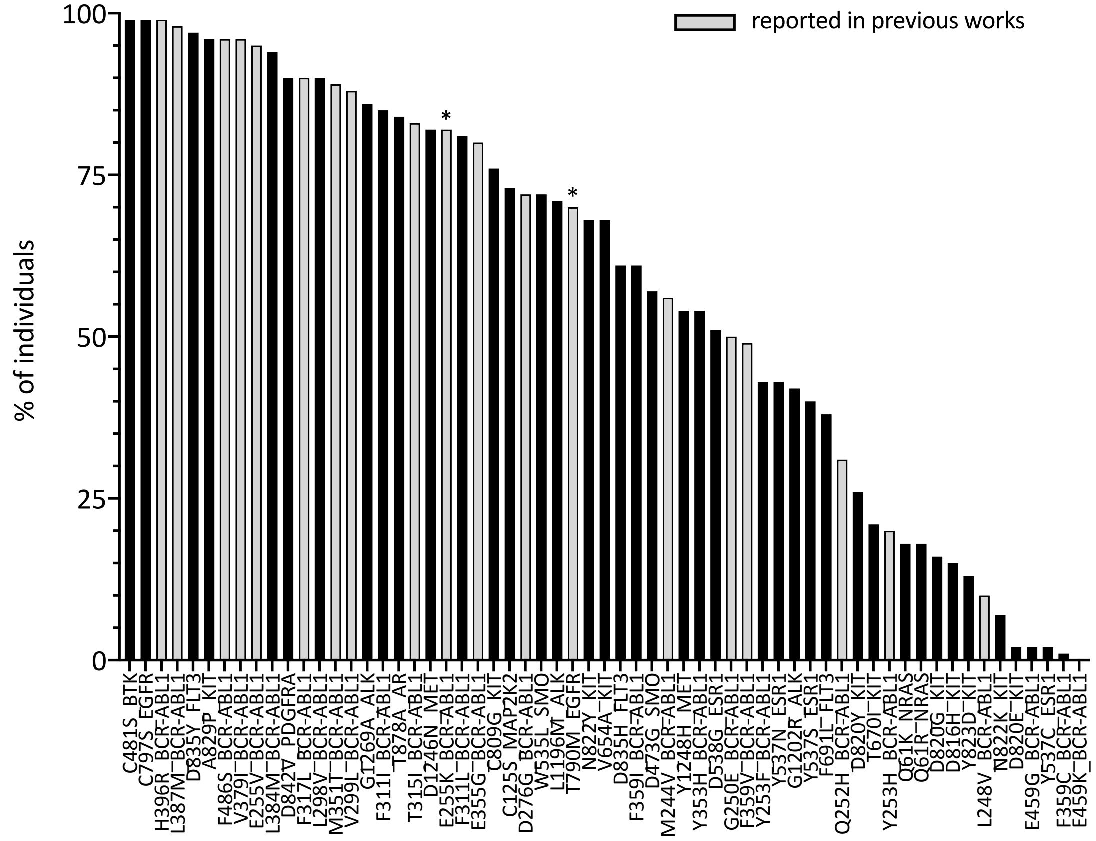


**Supplementary Figure 8.** *Same as Figure 3 with the only difference that instead of coloring the histogram's bars according to the tumor tissues the mutations occur in, we highlight in light grey what (to our knowledge) are all mutations that have been studied* in silico *in previous works for their likelihood to generate neopeptides that are HLA-presented* [1] [2] [3]*.* As in Figure 3, asterisks indicate what mutations in those works have further been tested *in vitro* and successfully shown to be able to elicit T-cell responses. Note that absence of asterisks from most previously studied mutations does not indicate that they cannot elicit T-cell responses but rather that they are not known to have been tested for it.

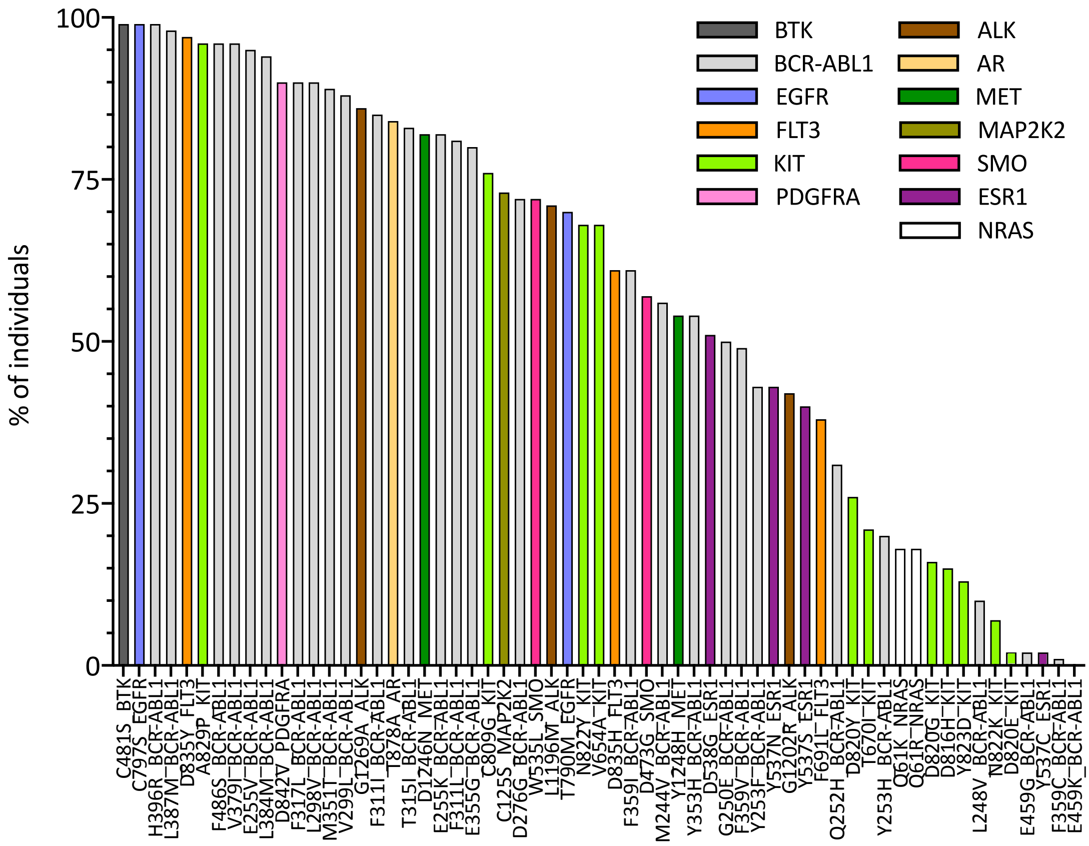


**Supplementary Figure 9.** *Same as Figure 3 with the only difference that histogram bars' colors indicate the different genes in which the mutations are found rather than the tumor tissues the mutations occur in.*

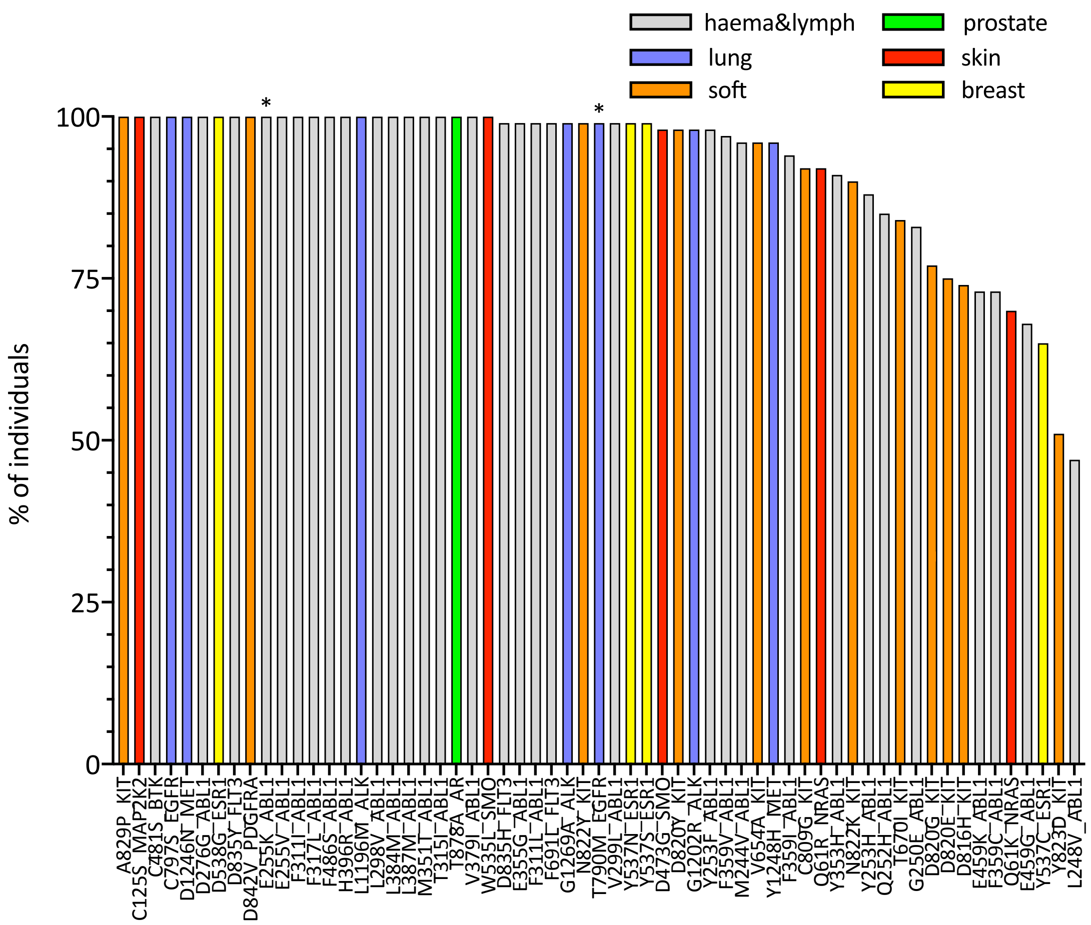


**Supplementary Figure 10.** *Equivalent of Figure 3 when considering a less restrictive patient-BR threshold below which individuals present a mutation's (i.e., patient-BR<2.0).*

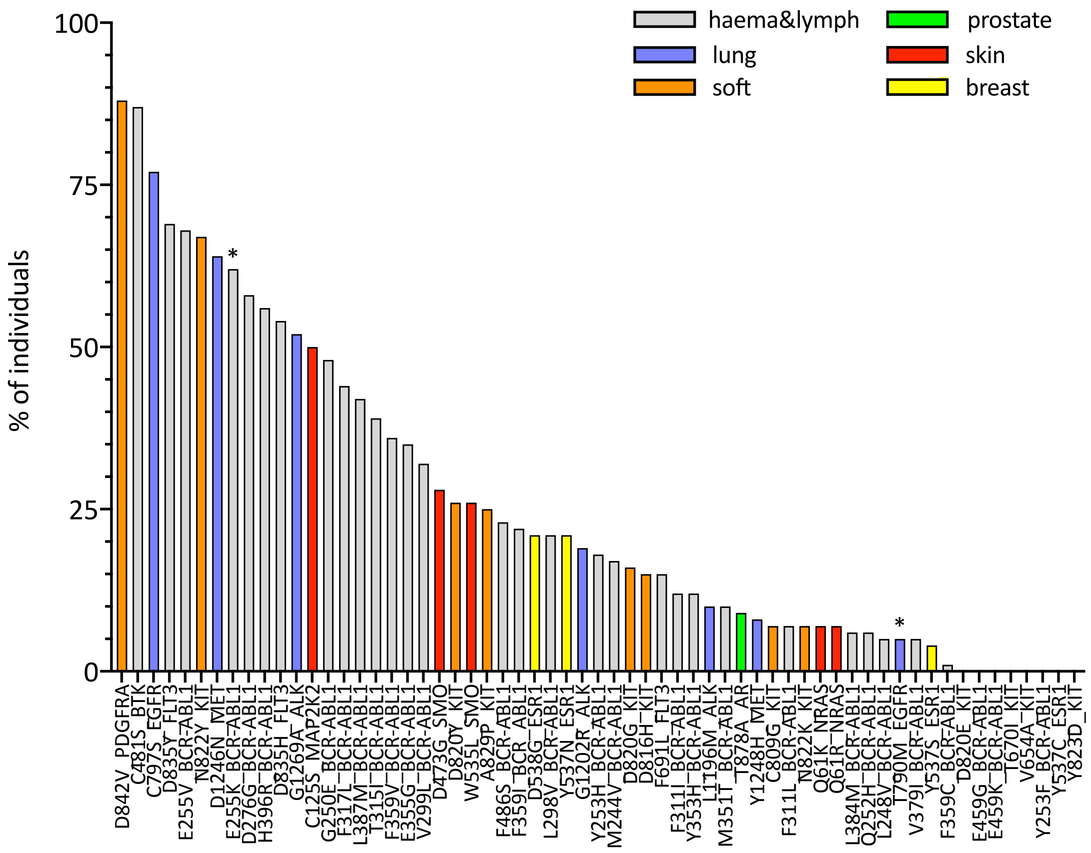


**Supplementary Figure 11.** *Equivalent to Figure 4 when using only HLA-A and HLA-B for calculating the patient-BRs (Equation 1).*

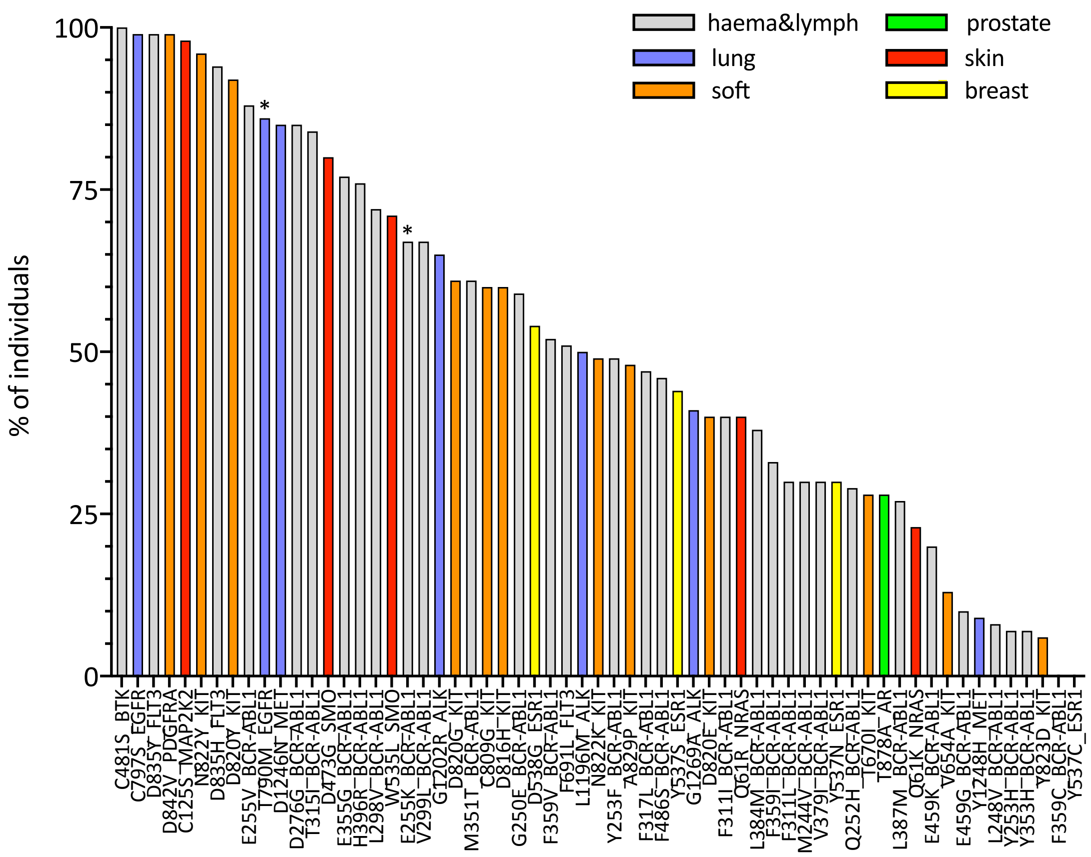


**Supplementary Figure 12.** *Similar to Figure 4 but using 2.0 as elution rank threshold.* This means that, for each mutation, the histogram illustrates the estimated percentage of individuals for which there exists at least one mutant-wild type peptide pair such that minimum eluted ligand likelihood percentile rank score across all of the patient's HLA types is <2.0 for the mutant peptide and >2.0 for the wild type peptide. Mutations on the x-axis are ordered according to decreasing percentages of individuals. We plot only mutations that have been observed in at least 5 patients (according to COSMIC). Colours indicate the different tumour tissues in which the resistance mutations have been observed; "haema&lymph" stands for haematopoietic and lymphoid tissue. Asterisks mark mutations that have been shown to elicit T-cell responses in previous works.

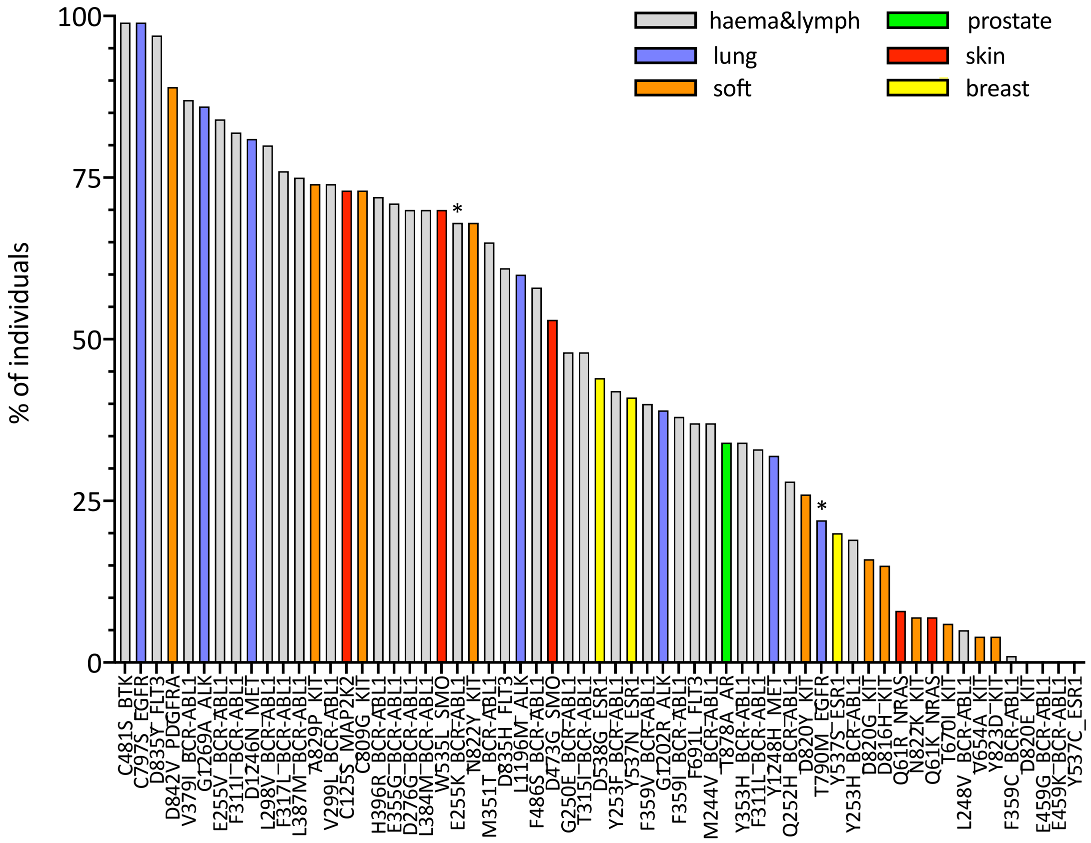


**Supplementary Figure 13.** *Similar to Figure 4 but asking that mutant peptides have eluted ligand likelihood percentile rank score <0.5 and lower than corresponding wild type peptides.* In other words, for each mutation, the histogram illustrates the estimated percentage of individuals for which there exists at least one mutant peptide-wild type peptide pair such that minimum eluted ligand likelihood percentile rank score across all of the patient's HLA types is <0.5 for the mutant peptide and for the wild type peptide is higher than for the mutant peptide (note: could still be <0.5). Mutations on the x-axis are ordered according to decreasing percentages of individuals. We plot only mutations that have been observed in at least 5 patients (according to COSMIC). Colours indicate the different tumour tissues in which the resistance mutations have been observed; "haema&lymph" stands for haematopoietic and lymphoid tissue. Asterisks mark mutations that have been shown to elicit T-cell responses in previous works.

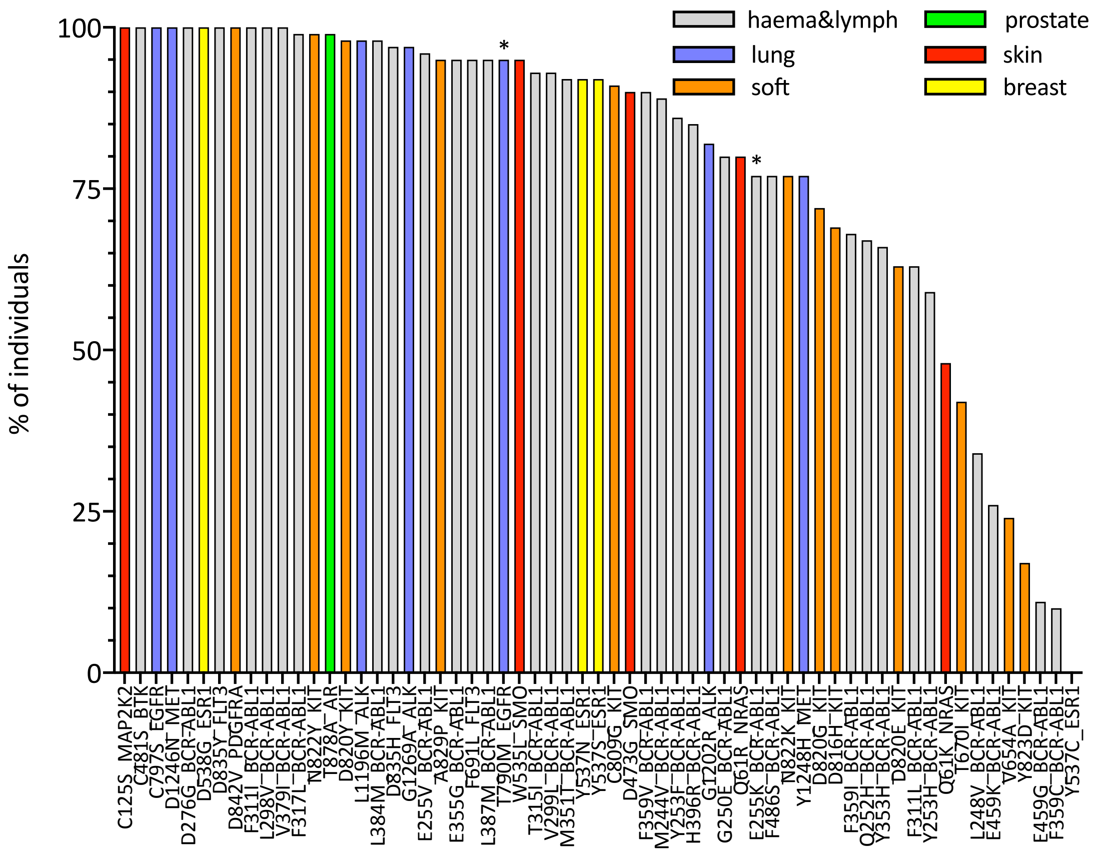


**Supplementary Figure 14.** *Similar to Supplementary Figure 13 but using an elution rank threshold of 2.0 instead of 0.5. .* In other words, for each mutation, the histogram illustrates the estimated percentage of individuals for which there exists at least one mutant peptide-wild type peptide pair such that minimum eluted ligand likelihood percentile rank score across all of the patient's HLA types is <2.0 for the mutant peptide and for the wild type peptide is higher than for the mutant peptide (note: could still be <2.0). Mutations on the x-axis are ordered according to decreasing percentages of individuals. We plot only mutations that have been observed in at least 5 patients (according to COSMIC). Colours indicate the different tumour tissues in which the resistance mutations have been observed; "haema&lymph" stands for haematopoietic and lymphoid tissue. Asterisks mark mutations that have been shown to elicit T-cell responses in previous works.

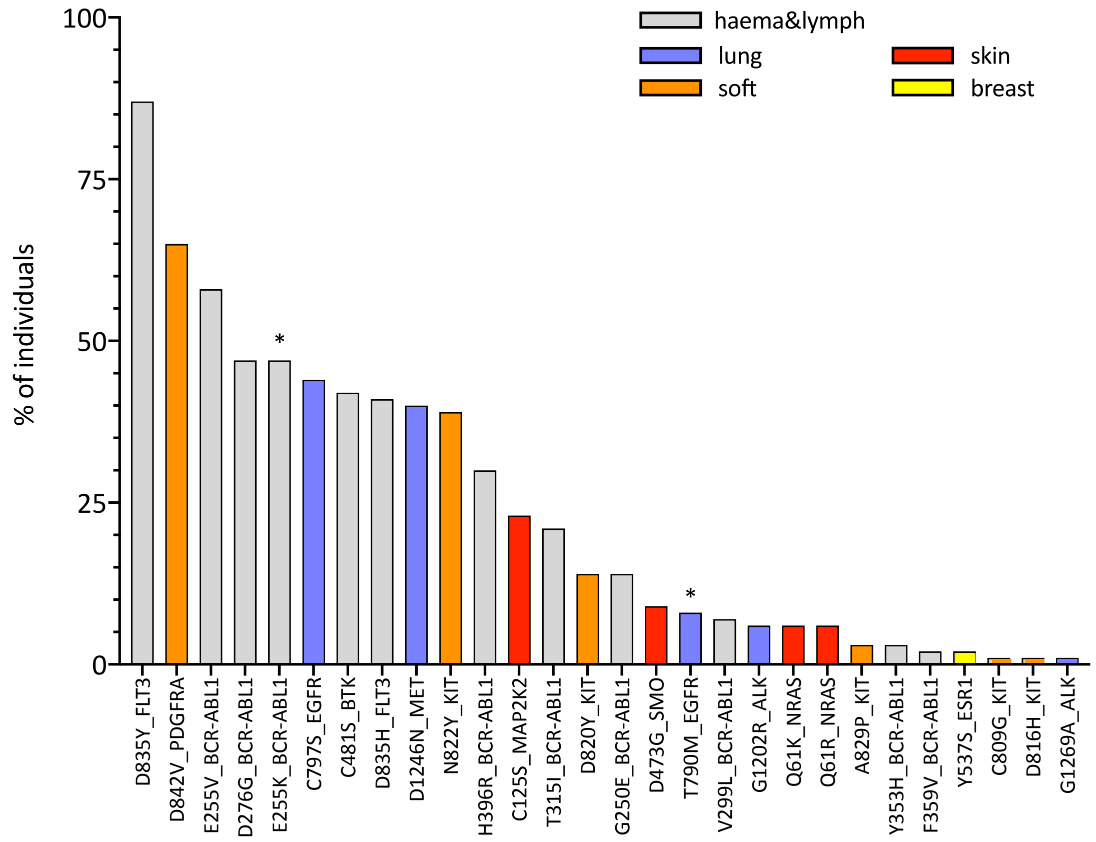


**Supplementary Figure 15.** *Similar to Figure 4 but asking that mutant peptides have eluted ligand likelihood percentile rank score <0.5 and corresponding wild-type peptides have eluted ligand likelihood percentile rank score >2.0.* This means that, for each mutation, the histogram illustrates the estimated percentage of individuals for which there exists at least one mutant-wild type peptide pair such that minimum eluted ligand likelihood percentile rank score across all of the patient's HLA types is <0.5 for the mutant peptide and >2.0 for the wild type peptide. Mutations on the x-axis are ordered according to decreasing percentages of individuals. We plot only mutations that have been observed in at least 5 patients (according to COSMIC) and for which the percentage of individuals is at least 1. Colours indicate the different tumour tissues in which the resistance mutations have been observed; "haema&lymph" stands for haematopoietic and lymphoid tissue. Asterisks mark mutations that have been shown to elicit T-cell responses in previous works.

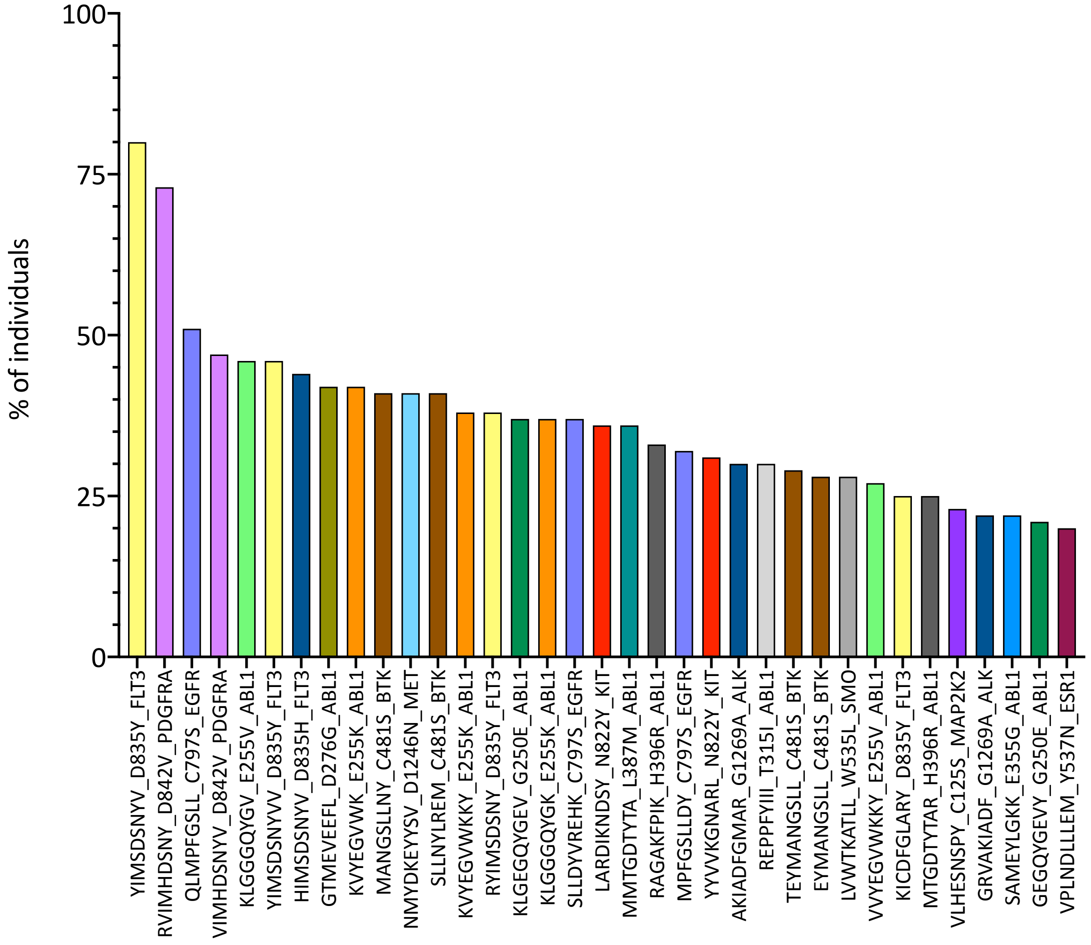


**Supplementary Figure 16.** *Estimates of the percentage of individuals in the general population for which a specific mutant peptide is predicted more likely to be presented with respect to its wild-type counterpart.* More specifically, the histogram illustrates the estimated percentage of individuals in which a given mutant peptide is associated to an eluted ligand likelihood percentile rank score across all of the individual's HLA allotypes <0.5 while the percentile score for its corresponding wild type peptide is >0.5. Mutant peptides on the x-axis are ordered according to decreasing percentages of individuals. We consider only mutant peptides from resistance mutations that occur in at least 5 patients in COSMIC and, for clarity, we plot only mutant peptides with the percentage of individuals satisfying the above conditions being >=20. Colours indicate peptides associated to different mutations.

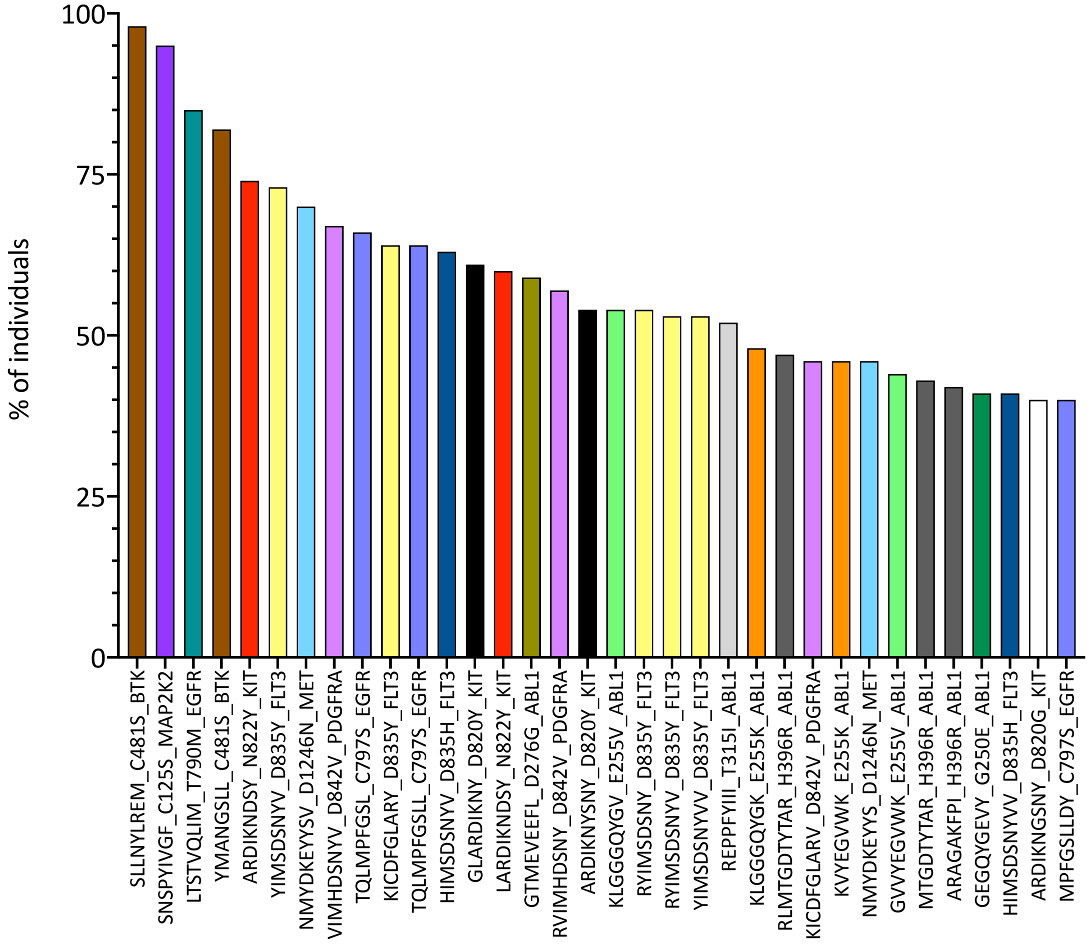


**Supplementary Figure 17.** *Similar to Supplementary Figure 16 but using an elution rank threshold of 2.0 instead of 0.5.* More specifically, the histogram illustrates the estimated percentage of individuals in which a given mutant peptide is associated to an eluted ligand likelihood percentile rank score across all of the individual's HLA allotypes <2.0 while the percentile score for its corresponding wild type peptide is >2.0. Mutant peptides on the x-axis are ordered according to decreasing percentages of individuals. We consider only mutant peptides from resistance mutations that occur in at least 5 patients in COSMIC and, for clarity, we plot only mutant peptides with the percentage of individuals satisfying the above conditions being >=40. Colours indicate peptides associated to different mutations.

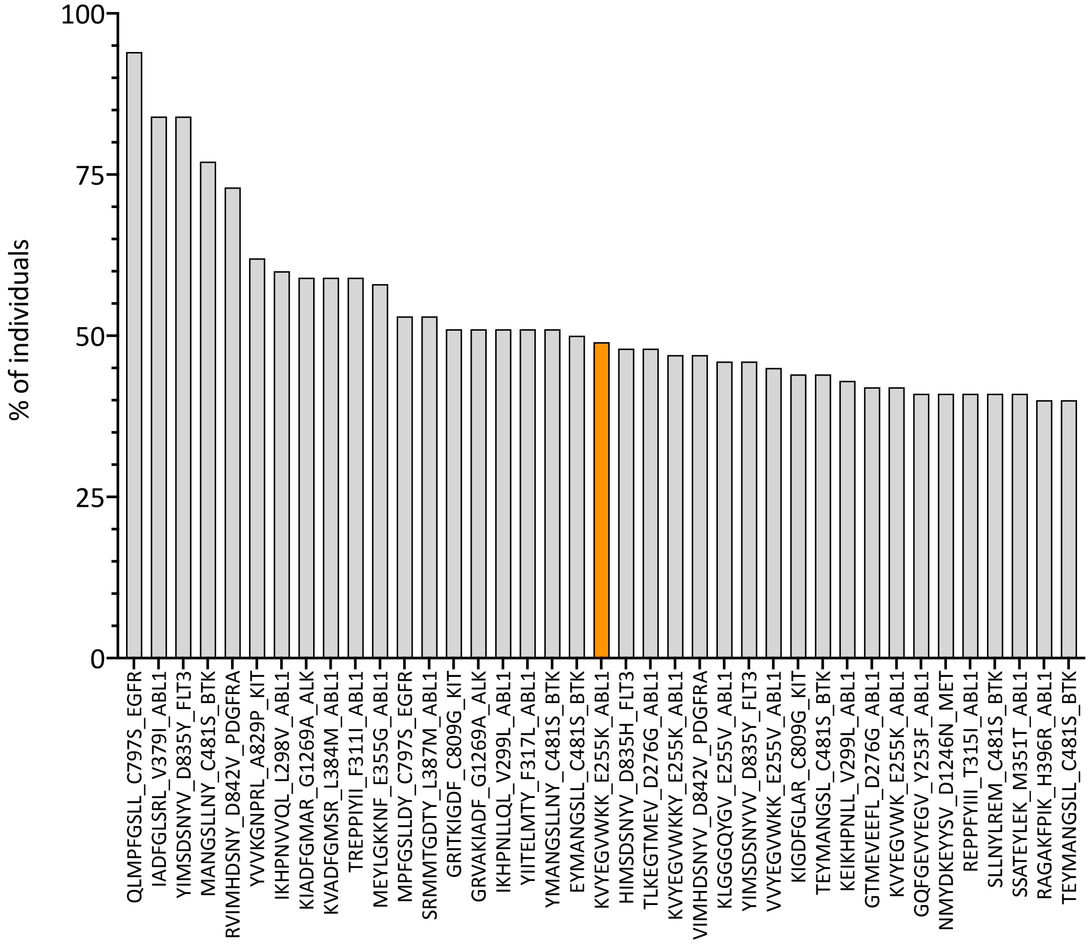


**Supplementary Figure 18.** *Similar to Supplementary Figure 16 but asking that wild-type peptides have simply higher elution rank than corresponding mutant peptides.* More specifically, the histogram illustrates the estimated percentage of individuals in which a given mutant peptide is associated to an eluted ligand likelihood percentile rank score <0.5 (when considering all of the individual's HLA allotypes) while the percentile score for its corresponding wild type peptide higher (considering all of the individual's HLA allotypes). Mutant peptides on the x-axis are ordered according to decreasing percentages of individuals. We consider only mutant peptides from resistance mutations that occur in at least 5 patients in COSMIC and, for clarity, we plot only mutant peptides with the percentage of individuals satisfying the above conditions being >=40. The orange bar highlights one of the peptides that in a previous study was shown *in vitro* to be able to elicit T-cell responses [1].

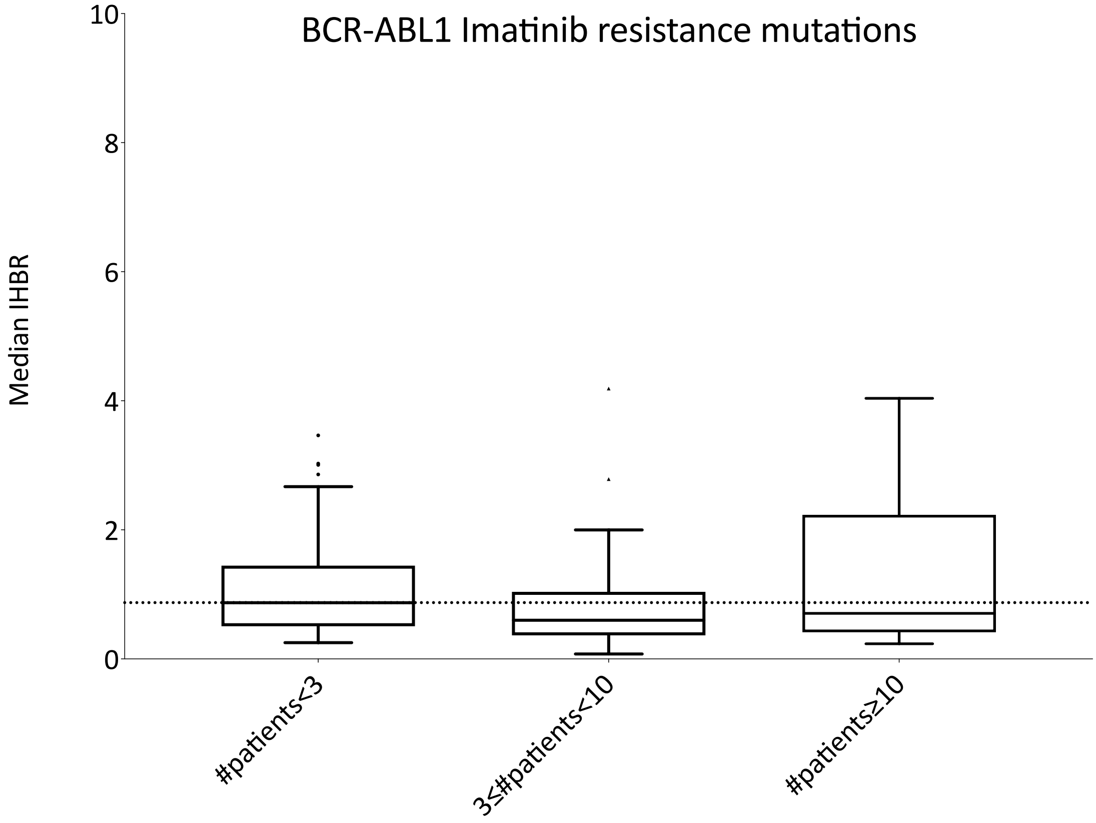


**Supplementary Figure 19.** *Distribution of PWHBR scores for 3 sets of BCR-ABL1 mutations that have been reported to confer resistance to the drug Imatinib. The different sets are characterised by a different number of patients in which the mutations have been observed (according to COSMIC).* Lower PWHBR values correspond to a higher likelihood of being presented by HLA class I complexes. The dotted horizontal line is a guide for the eye and corresponds to the value of the median of the distribution for mutations that have been observed in less than 3 patients. Note that while for consistency with previous graphs the y-axis is cut at 20, in this case no outlier point is found above this PWHBR value. The lower and higher edges of each Tukey box represent the 25% and 75% percentile value, respectively. The horizontal line inside each box represents the median value.

**REFERENCES.**


1 Cai, A. *et al.* Mutated BCR-ABL generates immunogenic T-cell epitopes in CML patients. *Clin Cancer Res* **18**, 5761-5772, doi:10.1158/1078-0432.CCR-12-1182 (2012).

2 Yamada, T. *et al.* EGFR T790M mutation as a possible target for immunotherapy; identification of HLA-A*0201-restricted T cell epitopes derived from the EGFR T790M mutation. *PLoS One* **8**, e78389, doi:10.1371/journal.pone.0078389 (2013).

3 Ofuji, K. *et al.* A peptide antigen derived from EGFR T790M is immunogenic in nonsmall cell lung cancer. *Int J Oncol* **46**, 497-504, doi:10.3892/ijo.2014.2787 (2015).
